# Chromosome-scale CRISPR screening reveals secretory pathway genes as drivers of aneuploidy-mediated antifungal tolerance

**DOI:** 10.64898/2026.07.23.740334

**Authors:** Nicholas C. Gervais, Brooke D. Esquivel, Lauren F. Wensing, Manini Chhina, Meea Fogal, Philippe C. Després, Lois L. Hoyer, Theodore C. White, Aleeza C. Gerstein, Rebecca S. Shapiro

## Abstract

The gain or loss of chromosomes in eukaryotes often drives aberrant phenotypes by altering the expression levels of hundreds or thousands of genes. In the case of beneficial aneuploidies, the genetic basis of fitness improvement has rarely been pinpointed, and identifying the causal genes remains a major challenge in engineering biology. The leading cause of human fungal infections, *Candida albicans*, frequently acquires extra copies of chromosome R (ChrR) following exposure to azole antifungal drugs, resulting in heightened antifungal tolerance. Despite this, why this aneuploidy frequently emerges following azole exposure and how it promotes antifungal tolerance remain unknown. Here, we combine RNA-seq with parallel chromosome-wide CRISPR activation (CRISPRa) and CRISPR interference (CRISPRi) screens to systematically profile the ChrR genes contributing to azole tolerance. Using multiplexed CRISPR-dCas12a tools, we further characterize the combinatorial effects of candidate genes and uncover a central role for post-Golgi secretory trafficking in antifungal tolerance. Specifically, we demonstrate that the secretory pathway regulators *SEC4* and *YPT31* are both necessary and sufficient for ChrR-mediated azole tolerance, and drive increased ergosterol content in the cell membrane. By leveraging a large-scale CRISPRa screen in a fungal pathogen, our work functionally dissects one of the most common aneuploidies observed in *C. albicans*, provides mechanistic insight into the molecular basis of antifungal tolerance, and establishes a generalizable framework for studying aneuploidy-mediated phenotypes across eukaryotic organisms.

## Introduction

Aneuploidy and large structural variations lead to an extraordinary diversity of functional outcomes in the cell^1^. While aneuploidy is almost always associated with adverse fitness consequences in normal human cells, it is observed ubiquitously in cancer and can drive phenotypes that are advantageous to cancer cells^2,3^. In the fungal kingdom, aneuploidy is common and is associated with a wide range of phenotypic consequences, including both fitness costs and adaptive responses to environmental stress^4,5^. Understanding the diverse ways by which aneuploidy shapes fungal evolution is critical, as fungi are a fundamental component of almost every ecosystem, and include numerous clinically important pathogens^6^. Human fungal pathogens cause over one billion infections annually, and the most common fungal pathogen, *Candida albicans*, is responsible for nearly one million life-threatening infections each year, with mortality rates exceeding 60%^7^.

The antifungal armamentarium available to treat invasive fungal infections is limited to only three major classes (azoles, echinocandins, and polyenes), and resistance is observed to all of them^8,9^. Aneuploidy is often associated with decreased antifungal drug susceptibility^4,8,9^, and the best-characterized example in *C. albicans* involves amplification of Chromosome 5 (Chr5) leading to resistance to the most widely used antifungal drug class, the azoles^10,11^. Because Chr5 encodes the azole drug target, Erg11, as well as a transcriptional regulator of efflux, Tac1, the corresponding upregulation of these genes via increased copy number is thought to underlie azole resistance^10,11^. Since the expression of thousands of genes is altered simultaneously in aneuploid fungal cells, it is very difficult to dissect which gene(s) underlie other aneuploidy-mediated drug susceptibility phenotypes^12–14^.

One of the most commonly observed aneuploidies following *in vitro* azole exposure is the acquisition of an extra copy of Chromosome R (ChrR). Previous work reports ChrR trisomies occurring between 50% and 100% of the time in response to azole antifungals^15–20^. Aneuploidies are often unstable and may be underreported in patient-derived strains^21^, though increased ChrR copy number has been detected in numerous clinical isolates, many following patient azole treatment^22–24^. Notably, ChrR trisomies are almost always associated with the emerging phenomenon of antifungal drug tolerance, rather than resistance. Antifungal tolerance refers to the ability of a subpopulation of fungal cells to grow slowly above the minimum inhibitory concentration (MIC) in a concentration-independent manner. Tolerance has been linked to persistent *C. albicans* infections and those that are recalcitrant to treatment despite being classified as susceptible by traditional clinical tests^9,25^. Unlike resistance, the exact genetic programs governing drug tolerance remain unclear, though they appear to be markedly diverse and polygenic in nature^9^. Taken together, there is strong evidence that ChrR aneuploidy is a fundamental mechanism by which *C. albicans* develops antifungal tolerance.

There are very few systematic approaches for dissecting the aneuploidy-phenotype relationship^26^, and developing scalable CRISPR technologies in fungi comes with unique challenges^27^. As aneuploidy-mediated phenotypes often result from the overexpression of multiple genes^10,28,29^, it is possible that overexpression of one or more genes on ChrR leads to heightened drug tolerance. Here, we combine whole-transcriptome analysis, parallel chromosome-wide CRISPRi/a-dCas9 screening, and multiplexed CRISPR-dCas12 approaches to precisely dissect the genetic basis for ChrR-mediated drug tolerance in *C. albicans*. Using this approach, we identify several novel regulators of antifungal tolerance. In particular, the simultaneous overexpression of two genes in the secretory pathway, *SEC4* and *YPT31*, was necessary and sufficient to fully recapitulate the tolerance observed in the ChrR trisomic strains. While ChrR trisomy reduced azole import, this effect was independent of secretory pathway expression and therefore did not contribute to the associated tolerance phenotype. Instead, upregulation of the secretory pathway increased ergosterol content in the cell membrane, providing a potential biochemical basis for secretory pathway-mediated azole tolerance. Together, we propose a new mechanism for one of the most frequently observed aneuploidies in *C. albicans*, and reveal an important pathway underlying the poorly studied phenomenon of antifungal tolerance in the leading cause of human fungal infections. This work establishes a pipeline to perform large-scale CRISPRa screens in fungal pathogens and to rapidly characterize aneuploidy-mediated phenotypes, offering a template that could be adapted to other organisms.

## Results

### ChrR aneuploidy led to distinct genetic reprogramming during growth in fluconazole

Given the high prevalence of ChrR trisomies in azole-tolerant *C. albicans,* we first aimed to assess the global transcriptional impact of ChrR trisomies via RNA-seq. We performed RNA-seq with a euploid *C. albicans* strain (SC5314) along with two ChrR-trisomic strains from the same strain background, cultured in nutrient-rich media (YPD) or in YPD supplemented with 64 µg/mL fluconazole (FLZ; **FIGURE 1A**). The two ChrR-trisomic strains differed in their haplotype of the extra copy of ChrR; one had an extra copy of the “A” haplotype ChrR (‘ChrR AAB’), and the other had an extra copy of the “B” haplotype (‘ChrR ABB’). We selected a high concentration of FLZ (64 µg/mL), well above the MIC (∼0.25 µg/mL), to characterize the transcriptional state of antifungal tolerance (slow growth above the MIC) rather than resistance.

**FIGURE 1:**
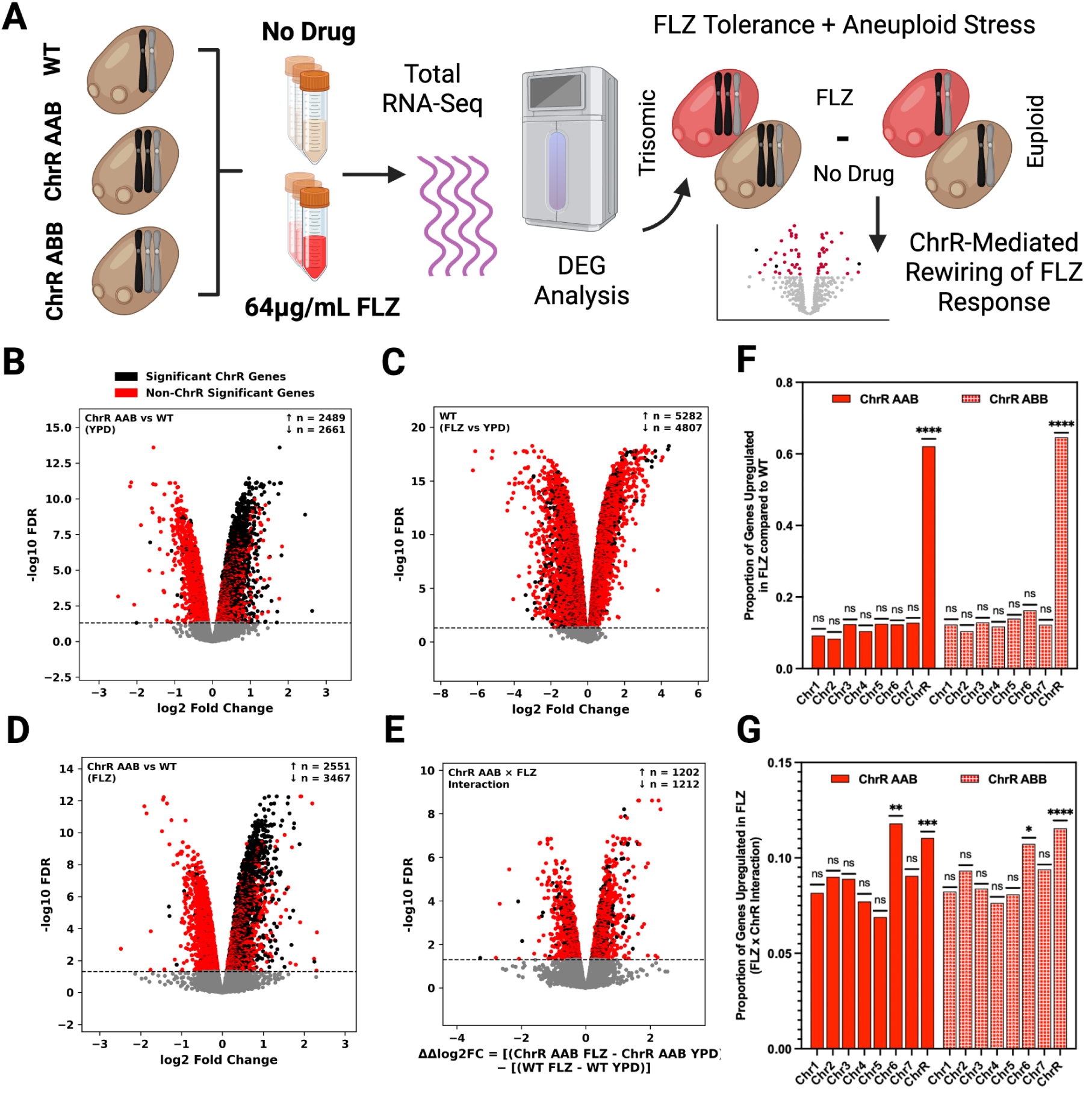
Chromosome R trisomies alter the transcriptional response to fluconazole. **(A)** Schematic of the RNA-seq experiment. Global gene expression changes in **(B)** one of the ChrR-trisomic strains (ChrR AAB) compared to the euploid WT in YPD **(C)** the euploid WT grown in 64 µg/mL FLZ compared to YPD **(D)** ChrR AAB compared to the euploid WT grown in 64 µg/mL FLZ without normalizing by DNA copy number, and **(E)** the aneuploid-conditioned FLZ response, i.e. the interaction between the ChrR trisomy and FLZ stress response in ChrR AAB (See Methods). *P*-values were adjusted for multiple testing using the Benjamini–Hochberg procedure. Genes (alleles) with FDR < 0.05 are coloured. **(F)** The proportion of genes by chromosome upregulated during growth in 64 µg/mL FLZ in both ChrR-trisomic strains (ChrR AAB and ChrR ABB) compared to the euploid WT, and **(G)** the proportion of genes by chromosome upregulated in the aneuploid-conditioned FLZ response in both ChrR-trisomic strains. Significance was tested for each chromosome using a hypergeometric test, ns *P* > 0.05 \**P* < 0.05, \*\**P* < 0.01, \*\*\**P* < 0.001, \*\*\*\**P* < .0001. Created in BioRender. Shapiro, R. (2026) https://BioRender.com/9qbjkmy.

The ChrR trisomy significantly reduced initial growth compared to the euploid strain (**FIGURE S1A**). Both drug treatment (PERMANOVA, R² = 0.73, *P* = 0.001) and genotype (R² = 0.13, *P* = 0.001) contributed significantly to global transcriptional variance (**FIGURES 1BC**, **FIGURES S1B**, **FIGURE S2**). Overall, we found that the expression of ChrR genes was ∼1.3-fold higher compared to non-ChrR genes in the ChrR-trisomic strains (**FIGURE S1C**), which is similar to previous findings for trisomies in *C. albicans*^12^. Of the genes that were the most highly upregulated in FLZ in all three strains (>3 log2 fold change [FC]), there was a significant enrichment of ChrR genes (hypergeometric tests: WT *P* = 0.008, ChrR AAB *P* = 0.013, ChrR ABB *P* = 0.010), suggesting FLZ caused a disproportionately strong upregulation of a handful of ChrR genes, regardless of ploidy (**FIGURES S2C-E**, **TABLE S1**). Both ChrR-trisomic strains exhibited significant transcriptional differences compared to the euploid strain when all strains were grown in FLZ (not normalized by DNA copy number) (**FIGURE 1D, FIGURE S2G**), where ChrR was the only chromosome significantly enriched for upregulated genes (∼63% of ChrR genes were upregulated) (**FIGURE 1F**).

We next sought to understand whether changes in gene expression in the ChrR-trisomic strains during growth in FLZ were simply due to their increased copy number, or a specific interaction between aneuploidy and drug. To account for the basal effect of the ChrR trisomy on gene expression, we subtracted the expression differences between the ChrR-trisomic strains and the WT during growth in YPD from the corresponding differences in FLZ (see Methods), yielding the ‘aneuploid-conditioned FLZ response’, which captures transcriptional changes beyond those attributable to increased ChrR copy number (**FIGURE 1EG**, **FIGURE S2H**). While the number of ChrR genes remained significantly enriched in the aneuploid-conditioned FLZ response, the proportion of upregulated ChrR genes was higher than other chromosomes by only ∼0.02 (**FIGURE 1G**). Upregulated genes on Chr6 and downregulated genes on Chr5 were also significantly enriched in the aneuploid-conditioned FLZ response, suggesting a potential relationship between ChrR and genes on other chromosomes during azole stress (**FIGURE 1G**) (**TABLE S1**). Indeed, ChrR and Chr6 trisomies frequently co-occur during adaptation to high concentrations of azoles^15,17,20^. Beyond these chromosome-level relationships, GO analysis revealed that several tRNAs were strongly repressed in the aneuploid-conditioned FLZ response, suggesting a possible role for mRNA translation regulation in the unique response of the ChrR-trisomic strains to stress (**TABLE S1**). Together, these findings suggest that ChrR-trisomic phenotypes arise from both preexisting changes in ChrR gene expression due to the trisomy and stress-specific changes in gene expression triggered by antifungal exposure.

### Parallel chromosome-wide CRISPRa and CRISPRi-dCas9 screening identified candidate genes for ChrR-mediated drug tolerance

Our RNA-seq analysis revealed widespread transcriptional alterations associated with ChrR trisomy, but it remained unclear which differentially expressed genes were responsible for the increased antifungal tolerance. Given that 994 genes reside on ChrR and many were upregulated in the trisomic strains, we next created parallel chromosome-wide CRISPRa and CRISPRi libraries to systematically identify genes contributing to antifungal tolerance (**FIGURE 2A**). For these ChrR-wide libraries, we exploited our previously developed *C. albicans*-optimized CRISPRa-dCas9^30^ and CRISPRi-dCas9^31^ systems for genetic overexpression and repression, respectively (rather than HyperdCas12a techniques), as they generally produce more modest changes in gene expression^32^, and therefore more closely approximate the expression differences observed in ChrR-trisomic strains. To maximize the likelihood of successful gene modulation, we designed up to seven sgRNAs targeting the promoter region of each ChrR ORF. Following filtering for predicted high-efficiency sgRNAs (see Methods), the final libraries contained 5,640 sgRNAs targeting 972 ChrR genes, including 780 genes targeted by at least six different sgRNAs (**TABLE S2**). Using an optimized pooled library construction pipeline^33,34^, we successfully generated *C. albicans* ChrR-wide CRISPRa and CRISPRi libraries with near-complete coverage, recovering >95% of the sgRNAs in the final library populations (**FIGURE S3**). Thus, we developed chromosome-scale CRISPR overexpression and repression libraries to enable the systematic dissection of aneuploidy-mediated antifungal tolerance.

**FIGURE 2:**
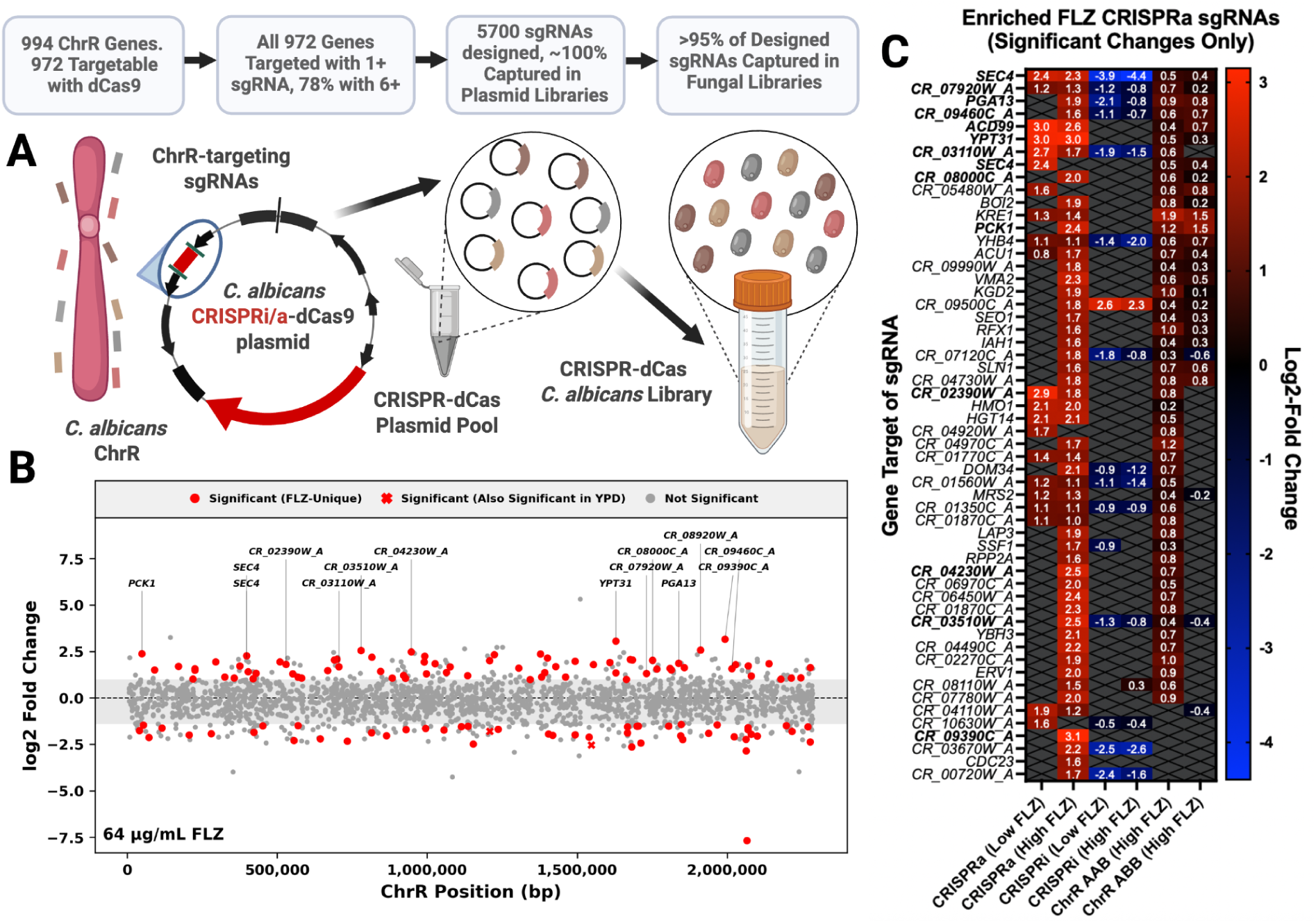
Parallel CRISPRi/a-dCas9 screens identify ChrR genes enriched or depleted in high concentrations of FLZ. **(A)** Schematic of the pooled ChrR CRISPRi/a-dCas library construction protocol with sgRNA design and capture statistics. **(B)** The changes in sgRNA abundance in the CRISPRa screen in 64 µg/mL FLZ. Each datapoint represents an sgRNA at its corresponding target position along the length of ChrR. The grey shaded area represents a 95% confidence interval based on the log2 fold-changes of the non-targeting sgRNAs. sgRNAs that were significantly enriched or depleted when the library was grown in plain YPD are marked with “X” since they likely do not represent changes specific to the 64 µg/mL FLZ environment. **(C)** Heat map of the sgRNAs that were enriched in the CRISPRa FLZ screens along with whether or not they were significantly enriched or depleted in FLZ in the CRISPRi screens, and whether or not they were up- or downregulated in one or both of the ChrR-trisomic strains in FLZ from the RNA-seq experiment. Only significant log2 fold-changes are displayed. Gene targets of sgRNAs that are bolded represent those sgRNAs that were selected for follow-up analysis. Created in BioRender. Shapiro, R. (2026) https://BioRender.com/gx7ws92.

To identify ChrR genes involved in mediating antifungal tolerance, we competitively passaged both CRISPR-dCas libraries in quadruplicate in YPD alone or supplemented with either 1 µg/mL or 64 µg/mL FLZ. We then used next-generation sequencing of the sgRNA region to compare the relative changes in strain abundance at the end of the experiment to the initial population. We selected a concentration near the MIC (1 µg/mL FLZ) to identify factors involved in both resistance and tolerance, and a concentration far greater than the MIC (64 µg/mL FLZ) to capture factors specifically leading to antifungal tolerance. The replicate screens had a strong positive correlation within each condition, supporting the overall quality and reproducibility of the screens (**FIGURE S4A**). Further, the 1 µg/mL FLZ and 64 µg/mL FLZ screens correlated positively with both CRISPRi/a systems, suggesting screening in the different FLZ concentrations yielded similar results overall (**FIGURES S4B-D**).

Both CRISPRa and CRISPRi screens revealed sgRNAs whose abundance changed specifically during growth in FLZ, representing candidate mediators of ChrR-associated antifungal tolerance (**FIGURE 2B, FIGURE S5**). To assess broad trends in the target genes corresponding to the set of 56 CRISPRa-enriched sgRNAs, we performed a STRING network analysis to identify protein-protein interactions (**FIGURE S6A**)^35^. While most proteins had no interactions, consistent with a lack of significant PPI enrichment (*P* = 0.533), a small module involving *CR_01750C* (*SEC4*), *CR_07520C* (*YPT31*), and *CR_01350C* (*SEC17*) formed (**FIGURE S6A**). This module involved the strongest overall interaction score in our analysis (0.94) between *SEC4* and *YPT31*, and the genes shared a role in the secretory pathway. Nearly all GO terms containing at least two genes from the 56 CRISPRa-enriched sgRNAs were also associated exclusively with *SEC4* and *YPT31* (**TABLE S1**). These observations provided preliminary evidence for the secretory pathway as an important mechanism in ChrR-mediated drug tolerance.

### Overexpression of individual ChrR genes conferred tolerance to azole antifungal drugs

To identify the strongest candidate drivers of ChrR-mediated antifungal tolerance, we selected 14 sgRNAs (representing 13 genes) from those that were the most enriched in the CRISPRa FLZ screens for follow-up study (**FIGURE 2C**, **TABLE S1**). We prioritized sgRNAs that were additionally upregulated in FLZ in the ChrR-trisomic strains from our RNA-seq data, as upregulation of genes on trisomic chromosomes often underlie aneuploidy-mediated phenotypes^10,28,29^. Selected sgRNAs that were also depleted in the FLZ CRISPRi screens provided further evidence that the corresponding target gene may be important for responding to FLZ stress (**FIGURE 2C**). These selected 14 sgRNAs were then used to generate independent CRISPRa strains.

We first used RT-qPCR to validate overexpression and found that 86% (12/14) of the resulting strains were indeed overexpressing their target gene (**FIGURES S6BC**). With the 12 successfully overexpressing CRISPRa strains, we measured FLZ resistance via IC_50_ changes at 24 h, and tolerance by calculating their supra-MIC growths (SMGs) at 48 h^9,25^. While none of the overexpression strains showed an increase in FLZ resistance, nearly all (10/12) showed a significant increase in FLZ tolerance, indicating that overexpression of these factors promotes azole tolerance rather than resistance, and confirming that the CRISPR screens were successful at selecting for tolerance, specifically (**FIGURE 3A**, **FIGURE S6D**). Several strains (7/12) also exhibited increased tolerance to posaconazole (POS), another azole to which ChrR-trisomic strains display enhanced tolerance^17^ (**FIGURES S6EF**).

**FIGURE 3:**
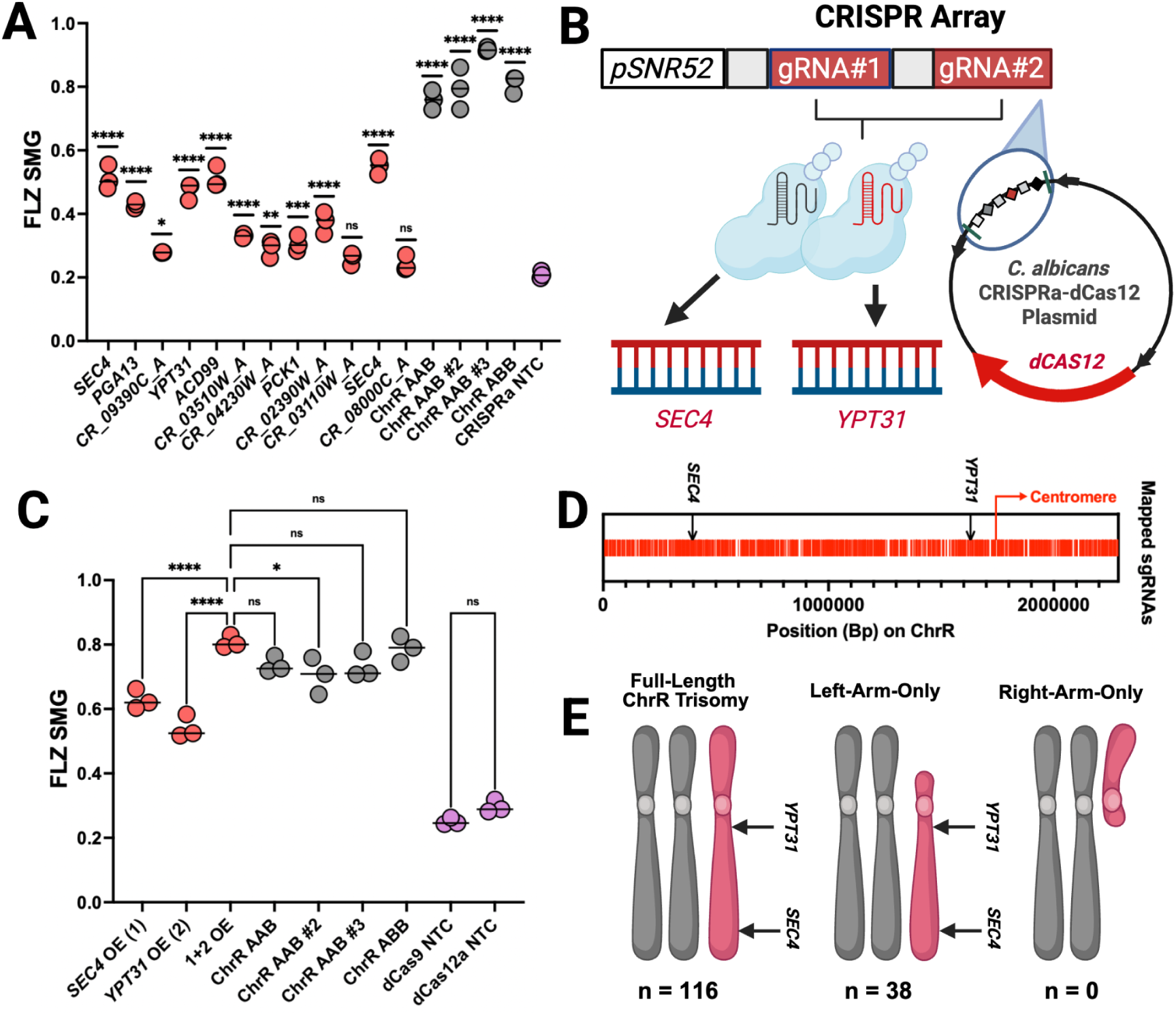
Upregulation of two secretory pathway factors is sufficient for ChrR-mediated antifungal tolerance. **(A)** The SMG (ie., average growth above the MIC at 48 h) for each strain grown in FLZ. Statistics were performed using an ordinary one-way ANOVA with Tukey’s multiple comparison test, where significance represents the difference between the given strain and the non-targeting control strain (NTC), ns *P* > 0.05, \*\**P* < 0.01, \*\*\**P* < 0.001, \*\*\*\**P* < 0.0001. Genes listed twice represent two distinct sgRNAs. **(B)** Conceptual schematic of the WT dCas12a-based multiplexed targeting of *SEC4* and *YPT31*. **(C)** The SMG of each strain in FLZ. Statistics were performed using an ordinary one-way ANOVA with Tukey’s multiple comparison test, ns > 0.05, \**P* < 0.05, \*\*\*\**P* < 0.0001. **(D)** The target position on ChrR of each designed sgRNA (red line) in the pooled libraries, with the location of *SEC4* and *YPT31* emphasized. **(E)** The frequency of ChrR trisomies in various strain backgrounds of *C. albicans* based on which arm(s) are triplicated following diverse *in vitro* selection strategies in different azole antifungal drugs^15–20^. *n* = number of independently-adapted populations the corresponding type of ChrR trisomy was observed in. Created in BioRender. Shapiro, R. (2026) https://BioRender.com/jzi7b5j.

Among these strains, three genes were responsible for the highest drug tolerance in both azole drugs (FLZ and POS) when overexpressed: *CR_01750C* (*SEC4*), *CR_08920W* (*ACD99*), and *CR_07520C* (*YPT31*) (**FIGURE 3A**, **FIGURE S6F**). Two different sgRNAs targeting *SEC4* were among the most enriched in the CRISPRa FLZ screens (**FIGURE 3A**, **FIGURE S5G**). To gain broader insight into the cellular functions of these top candidates, we assessed the growth of *SEC4*, *ACD99*, and *YPT31* mutants in response to diverse stressors, revealing that *YPT31* and *ACD99* shared responses to oxidative and hyperosmotic stress (**FIGURE S7**). Although little is known about the molecular function of *ACD99*^36^*, SEC4* and *YPT31* are both essential Rab family GTPases that regulate vesicle trafficking from the Golgi to the plasma membrane^37–39^. The identification of two secretory pathway components among the strongest tolerance-promoting hits further suggested that altered vesicle trafficking may play an important role in ChrR-mediated azole tolerance.

### Upregulation of the secretory pathway was sufficient to match ChrR-mediated azole tolerance

While overexpression of individual genes increased azole tolerance, none fully recapitulated the tolerance observed in four distinct ChrR trisomic strains (**FIGURE 3A**). This result suggested that ChrR-mediated tolerance likely arises from the combined effects of multiple genes on the chromosome. We therefore tested whether simultaneous overexpression of the strongest candidate genes (*SEC4*, *ACD99*, and *YPT31*) could more closely reproduce the trisomic phenotype, using a CRISPRa-dCas12a system previously optimized by our group for multiplexed gene regulation in *C. albicans*^32^. We selected crRNAs that led to a modest level of overexpression to best mimic trisomy-mediated increases in gene expression (**FIGURE S8A**). We then constructed CRISPR arrays targeting either all three of *SEC4*, *ACD99*, and *YPT31* together, or the two secretory pathway genes alone (*SEC4* and *YPT31*). The resulting strains overexpressed each target gene (**FIGURES S8B**, **S9AB**) to levels comparable to those observed in the ChrR-trisomic strains (**FIGURE S8C, FIGURE S9C**).

Strikingly, we found that the simultaneous overexpression of *SEC4* and *YPT31* was sufficient to recapitulate the elevated tolerance observed in all four independent ChrR-trisomic isolates (**FIGURE 3C**), and, like the ChrR-trisomic strains, did not alter FLZ resistance (**FIGURE S8D**). A similar pattern was observed with POS (**FIGURES S8EF**). The combined overexpression of *SEC4* and *YPT31* produced significantly greater azole tolerance than expected from their individual gene overexpression at the 24 h timepoint, based on a multiplicative model of genetic interaction (FLZ SMG; *P* = 0.008 and POS SMG; *P* = 0.008)^40,41^ (**TABLE S3**). In contrast, modest overexpression of *ACD99* did not measurably contribute to antifungal resistance or tolerance alone, or in combination with *SEC4* and *YPT31* (**FIGURE S9**). The combined requirement for increased gene dosage of both *SEC4* and *YPT31* is consistent with the frequent emergence of full-length and left-arm ChrR trisomies during azole stress^15–18,20^, as *SEC4* and *YPT31* reside near opposite ends of the left arm of ChrR (**FIGURE 3D**). Indeed, ChrR trisomies lacking amplification of the *SEC4* and *YPT31* loci essentially never occur in *C. albicans* following direct selection in various azoles (**FIGURE 3E**). Together, these findings identify the combined overexpression of two secretory pathway GTPases, *SEC4* and *YPT31*, as sufficient to explain the heightened azole tolerance conferred by ChrR trisomy.

### Upregulation of the secretory pathway was necessary for ChrR-mediated azole tolerance

We next asked whether elevated expression of *SEC4* and *YPT31* was required for the increased tolerance in the ChrR-trisomic strains, and thus whether restoring their expression to approximately euploid levels in a ChrR-trisomic background would restore azole susceptibility (**FIGURE 4A**). To test this, we used the CRISPRi-HyperdCas12a^32^ system to mildly repress both *SEC4* and *YPT31* in two ChrR-trisomic strains, thus mimicking the euploid level of expression of both genes in these otherwise trisomic strains (**FIGURE S8A; FIGURE S10A**). As aneuploid chromosomes may be unstable^42^, we validated that each of the trisomic CRISPRi strains had retained the ChrR trisomy and acquired no other changes in ploidy following the lithium acetate transformation (**FIGURE 4B**). Dual repression of *SEC4* and *YPT31* fully re-sensitized the SMG of both ChrR-trisomic strains to both FLZ and POS (**FIGURE 4C**, **FIGURE S10B**), without altering resistance (**FIGURE 4D**, **FIGURE S10C**). Therefore, *SEC4* and *YPT31* were both necessary and sufficient for ChrR-trisomy-mediated azole tolerance. By combining these gain- and loss-of-function approaches, we show that the adaptive phenotype conferred by an entire chromosomal trisomy can be explained by the altered dosage of just two genes.

**FIGURE 4:**
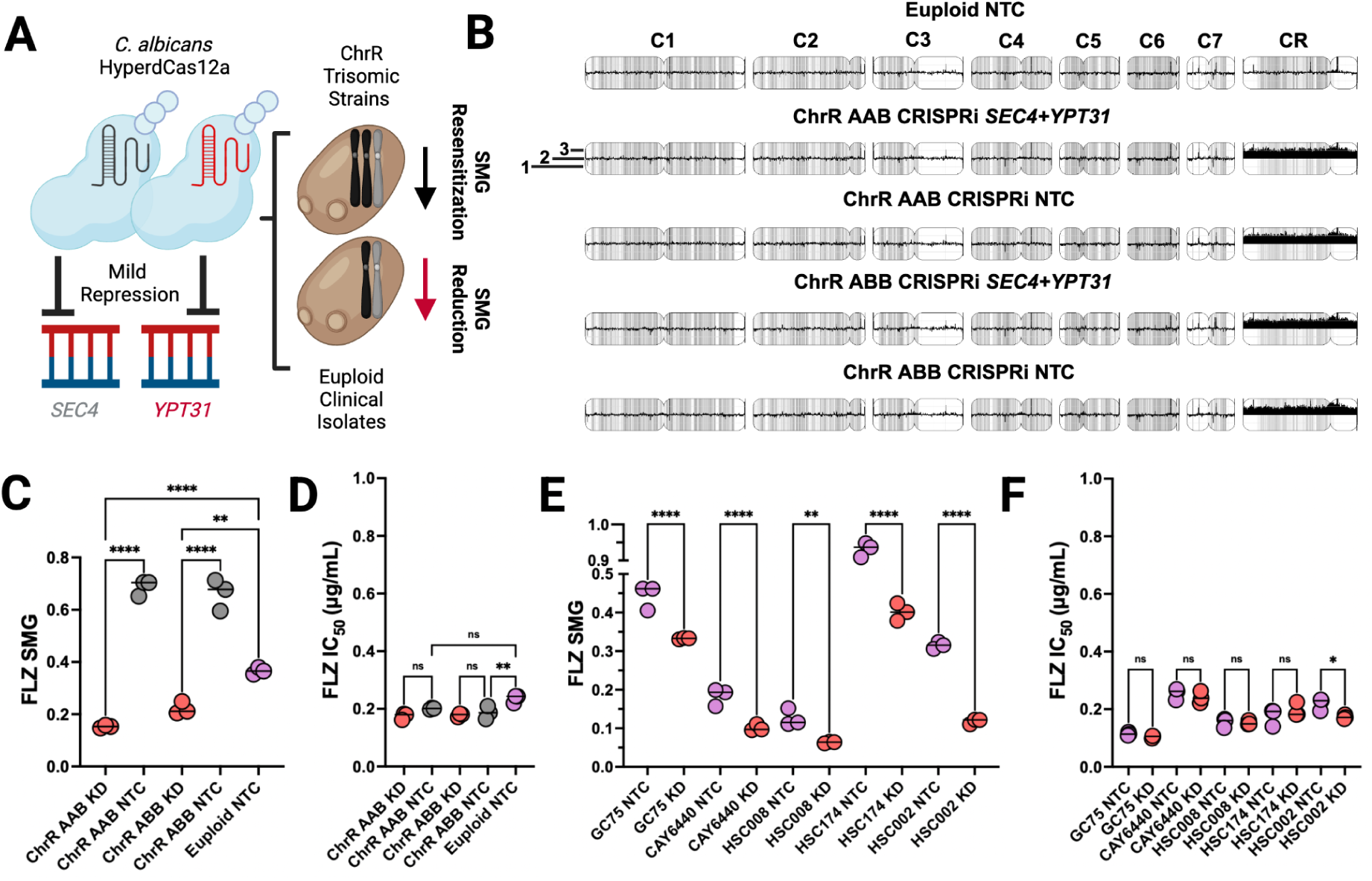
The secretory pathway represents a universal mechanism of azole antifungal tolerance which is necessary for ChrR-mediated antifungal tolerance. **(A)** Schematic for using CRISPRi-HyperdCas12a to return the expression of *SEC4* and *YPT31* in the ChrR-trisomic strains to euploid levels, and to slightly reduce the expression in clinical isolates. **(B)** Short-read sequencing data were uploaded to YMAP^44^ to confirm the expected copy number of ChrR in the CRISPRi ChrR-trisomic strains and absence of other structural variations. The copy number of each continuous DNA sequence along every chromosome (C1-CR) for all strains is labeled from 1-3 and plotted on the y-axis. **(C)** The SMG and **(D)** IC_50_ of each of the ChrR-trisomic *SEC4*+*YPT31* knockdown (KD) CRISPRi strains and the euploid non-targeting control (NTC) strain in FLZ. Statistics were performed using an ordinary one-way ANOVA with Tukey’s multiple comparison test, ns *P* > 0.05, \*\**P* < 0.01, \*\*\*\**P* < 0.0001. Comparisons may be statistically significant yet not meaningful based on standard antifungal susceptibility testing guidelines^43^. **(E)** The SMG and **(F)** IC_50_ in the *SEC4*+*YPT31* repression strain and corresponding CRISPRi non-targeting control strain (NTC) in five clinical isolate backgrounds. Statistics were performed using an ordinary one-way ANOVA with Šídák’s multiple comparisons test, ns *P* > 0.05, \**P* < 0.05, \*\**P* < 0.01, \*\*\*\**P* < 0.0001. Created in BioRender. Shapiro, R. (2027) https://BioRender.com/ro25vs5.

Mild repression of *SEC4* and *YPT31* in five distinct clinical isolates also significantly reduced SMG in both FLZ and POS with no meaningful changes in IC_50_ (**FIGURES 4EF**, **FIGURES S10DE**), given that the inherent variability of antifungal susceptibility testing necessitates a change of at least 2-fold for a change in IC_50_ to be significant^43^. Together, our findings establish secretory vesicle trafficking as a major and broadly conserved determinant of azole tolerance in *C. albicans* isolates, even in the absence of obvious chromosome size polymorphisms or large-scale karyotypic changes following growth in a high dose of FLZ (**FIGURES S10FG**), thus providing novel mechanistic insight into a clinically important phenotype whose molecular basis has remained poorly understood.

### Secretory-mediated tolerance drove increased membrane ergosterol and required drug transporters without altering efflux activity

*C. albicans* cells overexpressing efflux pumps accumulate increased concentrations of azoles in secretory vesicles^45^. To investigate the relationship between efflux and secretory-mediated antifungal tolerance, we overexpressed *SEC4* and *YPT31* via CRISPRa in a largely efflux-deficient (ED) strain (with *CDR1*, *CDR2*, *MDR1*, and *FLU1* deleted^46,47^), along with its corresponding reference strain (**FIGURE S11A**). The ED strains had a ∼4X decrease in FLZ resistance and had little to no FLZ tolerance compared to the parental strains, demonstrating the important role of efflux in antifungal tolerance (**FIGURE 5A**, **FIGURE S11BC**). While *SEC4*+*YPT31* overexpression still produced a small but significant increase in tolerance in the efflux-deficient background, the magnitude of this effect was greatly reduced relative to that observed in the reference strain (**FIGURE 5A**). These findings indicate that *SEC4*- and *YPT31*-mediated azole tolerance is largely dependent on drug efflux.

**FIGURE 5:**
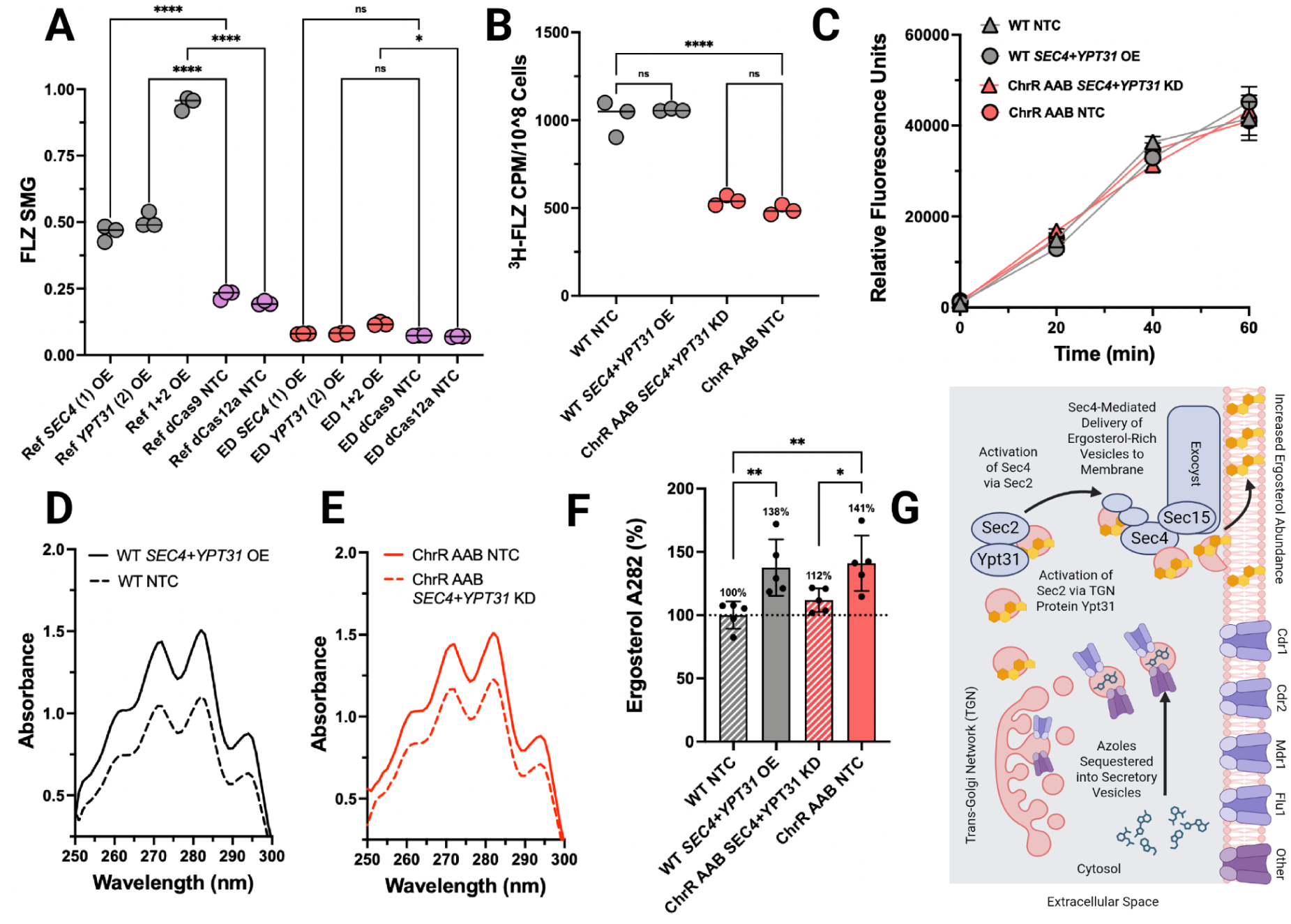
Secretory-mediated antifungal tolerance depends on drug transporters and drives increased membrane ergosterol. **(A)** The SMG values as a result of *SEC4* and/or *YPT31* overexpression in the reference strain (Ref) along with the derived efflux-deficient (ED) strain, all during growth in FLZ. Statistics were performed using an ordinary one-way ANOVA with Tukey’s multiple comparison test, ns *P* > 0.05, \**P* < 0.05, \*\*\*\**P* < 0.0001. **(B)** The number of radiolabeled fluconazole molecules (^3^H-FLZ) imported in the cell measured via counts per minute (CPM) per 10^8^ *C. albicans* cells. Statistics were performed using an ordinary one-way ANOVA with Šídák’s multiple comparisons test, ns *P* > 0.05, \*\*\*\**P* < 0.0001. **(C)** Extracellular Rhodamine 6G (R6G) fluorescence intensity over a 60-minute efflux period, measured in Relative Fluorescence Units (RFU). Error bars represent the standard deviation. **(D)** Absorbance spectra of sterols in the *SEC4*+*YPT31* overexpression strain compared to the euploid NTC, and **(E)** the ChrR-trisomic strain *SEC4*+*YPT31* knockdown (KD) strain compared to the ChrR-trisomic strain NTC. **(F)** The absorbance values at 282nm (representative of ergosterol abundance) for each strain reported as a % differential from the euploid WT NTC strain. Statistics were performed using an ordinary one-way ANOVA with Šídák’s multiple comparisons test, \**P* < 0.05, \*\**P* < 0.01. **(G)** Schematic of how the secretory pathway may interact with drug efflux and influence membrane ergosterol content towards azole tolerance. The trans-Golgi network protein Ypt31 activates Sec2, which activates and enables Sec4 to deliver secretory vesicles to the exocyst (via direct interaction with Sec15) for release from the cell^51^. In *C. albicans*, nascent drug transporters (including Cdr1, Cdr2, and Mdr1) embedded in the membrane of secretory vesicles pump intracellular azole molecules into the lumen of the vesicles^45^. Secretory pathway proteins could also facilitate the delivery of ergosterol-enriched secretory vesicles to the membrane^50^. Created in BioRender. Shapiro, R. (2027) https://BioRender.com/9xuv5xy.

Despite the involvement of efflux, we did not observe any relationship between intracellular radio-labeled fluconazole accumulation or R6G efflux and *SEC4*+*YPT31* expression (**FIGURE 5BC**). While we observed a decrease in fluconazole import with ChrR trisomy, it was independent of *SEC4+YPT31* expression and therefore unlikely to be a primary driver of ChrR-mediated azole tolerance (**FIGURE 5B**). The lack of change in efflux rate when altering *SEC4*+*YPT31* expression is consistent with the lack of a resistance phenotype following *SEC4*+*YPT31* overexpression. However, drug transporters pump intracellular azoles into secretory vesicles^45^, which can lead to decreased drug susceptibility in *C. albicans*^48^. Our data are therefore consistent with a model where efflux pumps on secretory vesicle membranes could facilitate the sequestering of azoles away from Erg11, as this would change the distribution of azoles in the cell without affecting overall accumulation.

As overexpression of *SEC4* and *YPT31* did not alter susceptibility to the ER stressor tunicamycin nor the ER-Golgi disruptor brefeldin A, the mechanism of secretory-mediated azole tolerance is unlikely to be a broad increase in secretory- or ER-based stress tolerance (**FIGURE S11DEF**). While ergosterol transport to the membrane may be primarily mediated by non-vesicular mechanisms^49^, increased secretory trafficking could enhance the delivery of ergosterol-enriched vesicles from the trans-Golgi network to the membrane^50^. Interestingly, we found that ChrR trisomy led to increased membrane ergosterol content, and that *SEC4+YPT31* overexpression was sufficient and necessary for driving this increase (**FIGURES 5DEF**). While a similar trend was observed, overexpressing *SEC4+YPT31* in the efflux-deficient strain did not lead to a significant change in ergosterol content (**FIGURE S11G**). Taken together, these data support the conclusion that secretory-mediated tolerance involves both a dependence on drug efflux and increased membrane ergosterol (**FIGURE 5G**).

## Discussion

While antifungal drug resistance mutations have been catalogued extensively in pathogenic fungi^52^, genes associated with tolerance are very poorly studied. Here, we characterize the relationship between ChrR aneuploidy and antifungal drug tolerance in *C. albicans*. This work represents a large-scale CRISPRa screen in a fungal pathogen, and highlights how previously inaccessible functional genetic questions are now addressable in *C. albicans* via the integration of CRISPR-based systems. Ultimately, we conclude that the secretory pathway genes *SEC4* and *YPT31* likely underlie ChrR-mediated drug tolerance in *C. albicans*. The essential nature of *SEC4* and *YPT31* highlights a critical advantage of these CRISPR-dCas tools, as other screening technologies that rely on gene mutation or deletion would likely fail to recapitulate these results. The reproducibility of these conclusions in several clinical *C. albicans* isolates suggests that this work may have revealed a mechanism of antifungal tolerance conserved across genetically diverse *C. albicans* isolates.

In addition to their role in the secretory pathway, Sec4 and Ypt31 are also required for autophagy in *S. cerevisiae*^53^. In *C. albicans*, azole tolerance may involve increased intracellular reactive oxygen species (ROS) leading to a reduction in membrane ergosterol autophagy^54^. Contrarily, other work suggests that the accumulation of ROS leads to loss of tolerance or unchanged levels of tolerance^55^. Here, we observed that *SEC4* overexpression leads to ROS resistance (**FIGURE S7C**), implying that autophagosome-related proteins would remain at the plasma membrane due to the relatively lower ROS burden, with correspondingly heightened rates of membrane autophagy^54^. Despite this, we observed increased membrane ergosterol upon overexpression of secretory pathway genes. ROS may therefore have context-dependent effects on azole tolerance. Taken together, our findings are consistent with a model where secretory-mediated azole tolerance could result in differential ergosterol degradation.

The strongest aneuploidy-mediated fitness increases usually involve the contributions of multiple genes^56^. Consistent with this model, the combined overexpression of *SEC4* and *YPT31* was necessary to match the level of antifungal tolerance in ChrR-trisomic strains. Our results therefore demonstrate how chromosome-scale copy number changes can generate phenotypes through coordinated regulation of multiple genes acting within the same biological pathway. The broader role that aneuploidy has in shaping tolerance at the population level, however, remains unclear. Future single-cell approaches could be used to dissect the role of aneuploid *C. albicans* cells in these heterogeneous drug tolerant populations.

While beneficial aneuploidies are associated with diverse phenotypes across the tree of life, the genetic basis of their fitness advantages remains largely unknown and represents a major obstacle in engineering biology^1^. Combining several new CRISPR-based strategies allowed us to identify the possible basis for one of the most commonly-observed aneuploidies in the leading cause of human fungal infections and its associated drug tolerance phenotype. It remains to be explored whether the secretory pathway also represents a key regulator of antifungal tolerance in diverse fungal pathogens. Together, this work provides a detailed mechanistic framework for how aneuploidy interacts with antifungal tolerance, thus helping to better understand the fundamental basis of antifungal tolerance, with future implications for the treatment of persistent *C. albicans* infections.

## Materials and Methods

### Plasmids, strains, and media

The plasmid backbones we used to perform our CRISPRi/a screens were our CRISPRa-dCas9-VPR (pRS156, Addgene #182707) and CRISPRi-dCas9-Mxi1 (pRS159, Addgene #122378) plasmids^30,31^. The plasmids we used to overexpress or repress combinations of genes were our constitutive CRISPRa-dCas12a (pRS1003, Addgene #247640), constitutive CRISPRi-HyperdCas12a (pRS1074, Addgene #247642), and inducible CRISPRi-HyperdCas12a (pRS1244, Addgene #247644) plasmids^32^. Information about all of the individual and multiplexed CRISPR-dCas *C. albicans* strains we made, including their corresponding gRNA sequence(s) and plasmid backbone, is available in **TABLE S4**.

The ChrR trisomic strains are from a previous experiment whereby *C. albicans* SC5314 was passaged a total of 10 times on yeast peptone dextrose (YPD)-agar plates supplemented with 1 µg/mL FLZ. At each passage, a single medium-sized colony was taken and struck onto a new plate, and the plate was incubated at 30°C for 48 h until the next transfer (manuscript in preparation). The efflux-deficient mutant used was DSY1024 (cdr1Δ::hisG/cdr1Δ::hisG cdr2Δ::hisG/cdr2Δ::hisG camdr1Δ::hisG/camdr1Δ::hisG flu1Δ::hisG/flu1Δ::hisG-URA3-hisG) along with its corresponding parental strain^46,47^.

*Candida albicans* strains were routinely cultured at 30°C (unless stated otherwise) in YPD, additionally supplemented with 250 µg/mL nourseothricin (NAT) from Jena Biosciences (cat. AB-102L) for plasmid selection when appropriate. 5-alpha Competent *Escherichia coli* cells from NEB (cat. C2987H) and 10-beta Electrocompetent E. coli from NEB (cat. C3020K) were grown at 30°C in Lysogeny Broth (LB) supplemented with both 100 µg/mL ampicillin (AMP) from BioShop (cat. AMP201.25) as well as 250 µg/mL NAT to help avoid plasmid recombination during selection. Fluconazole used for drug susceptibility screening and profiling was from Sigma-Aldrich (cat. PHR1160-1G), posaconazole was from Thermo Scientific Chemicals (cat. AC467660010).

### RNA extractions

Overnight yeast cultures were first diluted to an OD_600_ of 0.05 in 30 mL of YPD and left to grow at ∼240 rpm at 37°C for 4 h, whereby the OD_600_ values were all above 0.2. Cultures were then pelleted, resuspended in ∼1 mL of YPD, and frozen at −80°C. RNA extractions were performed using the PuroSPIN™ Total RNA Purification Plus Kit from Luna Nanotech (cat. NK251-200) with an enzymatic digestion using ∼300 μL of the thawed pelleted cultures. Briefly, pelleted cultures were resuspended in a freshly-prepared solution of 1 M sorbitol and 0.1 M EDTA adjusted to pH 7.4, 0.1% β-mercaptoethanol, and ∼200 units of Zymolase from BioShop (cat. ZYM001.1). The cell solutions were incubated at 30°C for approximately 45 min with gentle agitation, then centrifuged at 4000 × *g* for 5 min to obtain a pellet of spheroplasts. After the supernatants were discarded, 400μL of the Working Buffer LB-R (supplemented with 20μL of β-mercaptoethanol per 1mL) was added, and the spheroplasts were vortexed thoroughly for 1-2 min until they were completely resuspended. All subsequent steps were performed exactly as per the manufacturer’s instructions. RNA was quantified via NanoDrop using an Infinite 200 PRO microplate reader (Tecan).

### Aneuploid strain drug screening and RNA-sequencing

The strains tested with RNA-seq were the euploid WT (SC5314), strain ChrR AAB, and strain ChrR ABB. To compare basal expression levels in the euploid and ChrR-trisomic strains and assess their responses to FLZ treatment, one 5 mL YPD culture of each of the three strains was prepared and grown overnight at 30°C. The next morning, each strain was diluted to an OD_600_ of 0.05 in 3 tubes of 15 mL YPD, as well as 3 tubes of 15 mL YPD with 64 µg/mL FLZ, and left to grow at 37°C (∼240 rpm) for 6 h. Following this, the OD_600_ values were read, all 18 samples were pelleted at 4000 rpm for 5 min, resuspended in ∼1 mL of YPD, and frozen at -80°C. Differences in the OD_600_ values following the 6 h of growth were tested using a two-way ANOVA followed by a Šídák’s multiple comparisons test with GraphPad Prism version 10.5.0 for macOS, GraphPad Software, Boston, Massachusetts USA, www.graphpad.com.

RNA was extracted as described above and analyzed via TapeStation to confirm the samples were of high quality. Total RNA-seq library prep was performed by the Advanced Analysis Centre at the University of Guelph using a combination of the QIAseq FastSelect - rRNA Yeast Kit from Qiagen (cat. 334215) for rRNA depletion, followed by library prep with a NEBNext® Ultra™ II Directional RNA Library Prep Kit for Illumina® from NEB (cat. E7760L). Libraries were sequenced with the Illumina NovaSeq X platform by SeqCenter in Pittsburgh, PA. The raw sequencing files have been deposited to the SRA (BioProject ID: PRJNA1471677).

### RNA-seq analysis

Our pipeline for analyzing the RNA-seq data is available on our Github: https://github.com/TheShapiroLab/ChromosomeR. Reads were not initially trimmed to avoid introducing biases into differential expression estimates^57^. Paired-end reads were aligned to the reference *C. albicans* SC5314 genome, Assembly 22 downloaded from the *Candida* Genome Database (CGD)^36^, using STAR (2.7.11b)^58^. We output the alignments as coordinate-sorted .BAM files (--outSAMtype BAM SortedByCoordinate). To avoid spurious splice alignments, especially considering introns are rare in *C. albicans*^59^, non-canonical splice junctions were filtered out (--outFilterIntronMotifs RemoveNoncanonical). Because both homologous chromosome sequences are represented in the diploid reference genome we used for alignment, as well as the potential presence of paralogous or repetitive genomic regions, we allowed multimapping reads to align to multiple genomic loci, though a single alignment was then randomly selected and reported for downstream analysis to avoid counting a read twice (--outFilterMultimapNmax 100, --outSAMmultNmax 1, --outMultimapperOrder Random, --runRNGseed 42). Gene-level read counts and transcriptome-aligned .BAM files were generated during alignment (--quantMode GeneCounts TranscriptomeSAM). Alignment metadata tags were also generated to support downstream analyses (--outSAMattributes NH HI AS nM XS). Gene-level counts were then generated using featureCounts^60^, and differential expression was tested via edgeR^61^. All of the outputs were combined into a single .csv file, seen in **TABLE S1**.

To calculate the aneuploid-conditioned FLZ response in the ChrR-trisomic strains, we started with the log2 FCs of each gene in FLZ compared to YPD for both trisomic strains, thus providing us with information about how each gene differed in FLZ compared to YPD. We then subtracted the corresponding log2 FCs when the trisomic strains were compared to the WT during growth in plain YPD, thus accounting for basal differences in gene expression between the three strains. The resulting log2 FCs therefore represent how differentially expressed genes during growth in FLZ compare in each ChrR-trisomic strain to the WT.

### Design and synthesis of the sgRNA library

To design the sgRNA library, we first retrieved the flanking sequences for all *C. albicans* SC5314 ChrR genes (haplotype A) using the CGD Batch Download Tool^36^. We proceeded with only the ORF features (including dubious ORFs) as well as the four annotated non-coding RNAs on CGD. We have previously demonstrated that sgRNA success may depend on location relative to the transcriptional start site (TSS) in some cases, though current *C. albicans* TSS annotations do not always lead to improved sgRNA efficacies compared to simply targeting a region upstream of the start codon^30^. Additionally, TSSs are not annotated for every gene. Therefore, we designed sgRNAs in a different region of the promoter based on whether the given gene had an annotated TSS and how far the TSS was relative to the start codon. In every case, sgRNAs were designed 90-370 bp upstream of the start codon or TSS, as we have previously identified this to be a broadly successful range in which our CRISPR-dCas systems result in successful sgRNA activity^30–32^. For ORFs with a predicted dominant TSS of =< 140 bp, 6 sgRNAs were designed upstream of the start codon, and no additional guides targeting the TSS. 140 bp was selected as the cut-off since this is the last point at which at least half of the targeted range from 90-370 bp upstream will include the ideal target range of the TSS for all ORFs included in this group. For ORFs with a predicted dominant TSS of >140 bp, 3 sgRNAs were designed upstream of the start codon, with an additional 4 sgRNAs targeting upstream of the TSS. For ORFs with no predicted dominant TSS, 4 sgRNAs were designed upstream of the start codon, with an additional 3 sgRNAs targeting 370-670 bp upstream of the start codon to ensure sgRNAs exist upstream of the true dominant TSS if it is relatively far upstream. 570 bp was selected as the cut-off as 95% of known *C. albicans* TSSs are less than 480 bp upstream of the start codon. Thus, the additional 90 bp leaves room for sgRNAs designed upstream of them. For the ncRNAs targeted, up to 10 sgRNAs were designed in a range of 0-500 bp upstream of the feature start. Our code to isolate a corresponding region of the promoter for each gene before designing sgRNAs can be found on our Github: https://github.com/TheShapiroLab/Pooled-Library-Build.

After the corresponding promoter regions were isolated, we used the Eukaryotic Pathogen CRISPR guide RNA/DNA Design Tool (batch mode) to output all of the corresponding possible sgRNAs per gene (SpCas9: 20 nt gRNA, NGG PAM on 3’ end, *C. albicans* SC5314 FungiDB-26)^62^. sgRNAs were removed if they included any non-ATCG nucleotides (ie. “N” nucleotides from the imperfect *C. albicans* assembly), any perfect or imperfect off-target effects, both of a pair of duplicate sgRNA sequences (ie. if overlapping or neighbouring genes could be targeted successfully by the same sgRNA), or any with SapI or PacI RE sites as they would effectively be destroyed during downstream cloning and fungal library construction steps. After this, the corresponding number of sgRNAs were selected for each gene based on the total efficiency score. Note that in a few rare cases, different sgRNAs were given identical names as names provided by the Eukaryotic Pathogen CRISPR guide RNA/DNA Design Tool are derived from the target gene as well as the position of the sgRNA in the uploaded sequence, and there were multiple different target regions provided for many genes (ie. regions upstream of the start codon vs. TSS). The final N20 sgRNA sequences were flanked with sticky overhangs used for cloning along with SapI RE sites, as well as primer binding sites for library amplification following synthesis. For example: 5’-CTCTTGTCTGGCTACTTTCGG**GCTCTTC**ccgaTGGGAGGGACTTGTTCAATAgttg**GAAGAGC**C GAAACCAGCCATCAGCCAAG -3’, where bolded nucleotides correspond to the SapI RE sites, the lowercase nucleotides correspond to the sticky overhangs once treated with SapI, and the region in between corresponds to the N20 sgRNA sequence. Non-targeting sgRNAs (20 bp) were then adapted from a previous yeast CRISPRi screen^63^ and kept only if there were at least 5 mismatches to the *C. albicans* genome using the CGD BLASTn tool^36^, to a final proportion of ∼1% of the library, as is standard in other systems^63,64^. Non-targeting sgRNAs were additionally filtered for the same criteria as stated for the targeting sgRNAs, if relevant. The final library has 5700 sgRNAs. All sgRNA sequences can be seen in **TABLE S2**. The oligo pool was purchased from and synthesized by Twist Bioscience (South San Francisco, CA, USA).

### Mapping the sgRNAs to the genome

To map the sgRNAs in the oligo pool to their bp position on ChrR, we first downloaded the reference *C. albicans* SC5314 Assembly 22 genome from the CGD^36^ and isolated the sequence for ChrR. Next, we used Bowtie^65^ to map the sgRNA sequences to ChrR with the -f -y -a -m 1 -v 1 -x options, -m being important to prevent an sgRNA aligning to multiple genomic regions. Of the 5700 reads (sgRNAs) processed, 5640 (98.95%) aligned, 60 failed to align (1.05%), and 82 alignments were suppressed due to -m (1.44%). Initially, the PAM sites were included in the alignment to help improve specificity, though the output sgRNA positions were therefore off by 3 bp in situations where there was an sgRNA on the opposite strand of a Watson-strand gene (“revcom”), or when there was an sgRNA on the same strand as a Crick-strand gene (non-”revcom”). Therefore, these sgRNAs were increased by 3, and all sgRNAs regardless of orientation and strand were increased by 10 to reflect the middle position of the sgRNA. Positions of each sgRNA are listed alongside the screening data in **TABLE S1**.

### Plasmid library construction and verification

We first resuspended the sgRNA oligo library to a concentration of 10 ng/µL in pH 8.0 TE buffer, then performed a primer extension reaction using the KAPA HiFi HotStart plus dNTPs (100 U) kit from Roche (cat. no. 7958897001). Briefly, we mixed 10 ng of the pooled oligos, 0.75 μL each of the library amplification primers at 10 μM (see **TABLE S4**), 0.75 μL of dNTPs, 5 μL of 5X KAPA HiFi Fidelity Buffer, 0.5 μL of KAPA HotStart DNAP, and nuclease-free water to 25 μL. We next performed a PCR with the following conditions: 1) 3 min at 95°C, 2) 20 s at 98°C, 3) 15 s at 69°C, 4) 15 s at 72°C, steps 2-4 10 times, and 5) 1 min at 72°C. We then purified the PCR product using a DNA Clean & Concentrator-5 kit from Zymo Research (cat. no. D4013). Finally, we prepared a Golden Gate reaction mix containing ∼5000 ng of the receiving plasmid backbone, 2 μL of 10X rCutSmart Buffer, 2 μL of ATP from NEB (cat. P0756S), 1 μL of T4 DNA ligase from NEB (cat. M0202S), 1 μL of SapI from NEB (cat. R0569L), and nuclease-free water to 20 μL. Reactions were performed in a thermocycler using: 37°C for 2 min and 16°C for 5 min for 99 cycles, followed by 65°C for 15 min, and 80°C for 15 min. Following this, an additional 1 µL of SapI was added to each reaction and incubated at 37°C for 1 h to remove any uncloned plasmids.

Following this, the cloned product was transformed in duplicate into 10-beta Electrocompetent *Escherichia coli* cells from NEB (cat. C3020K). First, 40 μL of cells thawed on ice were mixed with 1.5 μL of the cloned plasmid pool. The entire volume was then transferred into one of the Electroporation Cuvettes Plus from Fisherbrand (cat. FB102), which had been left in the freezer to cool. Electroporations were performed with a MicroPulser^TM^ Electroporation Apparatus from BioRad with the ‘EC2’ preset (2500 V for 5.0 ms). Immediately afterward, 1 mL of SOC outgrowth medium was added directly to each cuvette, and all were left to recover at 37°C (∼240 rpm) for 1 h. Following this, a small aliquot of both reactions were plated onto one agar plate of LB + AMP + NAT each to confirm the rate of transformation efficiency, and left to incubate at 37°C static. The rest of the reactions were inoculated into one flask each of 500 mL LB + SeaPrep Agarose (at 0.35%) from Lonza (cat. 50302) with 100 µg/mL AMP and 250 µg/mL NAT. Both flasks were left to sit on ice for ∼1 h to allow the agar to partly solidify, and were then moved carefully to incubate at 37°C for ∼30 h until colonies were visible. After this, both flasks were combined and pelleted at 4696 x *g*, washed in fresh LB three times to remove residual agarose, and then mixed with 40% glycerol, aliquoted into separate tubes, and left at -80°C. The library was then struck out onto LB + AMP + NAT and the plasmids from individual colonies were sequenced to tentatively confirm successful library construction before next generation sequencing. The NucleoBond Xtra Maxi Plus EF kit from Macherey-Nagel (cat. 740424.1) was used to prep high concentrations of the plasmid library for fungal library construction as per the manufacturer’s instructions and as previously described^33,34^.

### Candida albicans transformation

*C. albicans* cells were grown up overnight in YPD at 25°C (∼240 rpm) to help ensure the cells were still in growth phase the following day. Plasmids were linearized using PacI from NEB (cat R0547L), and simultaneously digested with SapI from NEB (cat. R0569L) to help remove any remaining plasmids without a successfully cloned sgRNA. The double digests were left at 37°C for 12 h, followed by a 20 min heat-inactivation step at 65°C. A transformation master mix was prepared with 800 µL of a solution of 50% polyethylene glycol (PEG) from BioShop (cat. PEG335.1), 100 µL of 10X Tris-ethylenediaminetetraacetic acid (EDTA) buffer solution, 100 µL of a 1 M lithium acetate solution, 40 µL of UltraPure™ Salmon Sperm DNA Solution from ThermoFisher (cat. 15632-011), and 20 µL of a solution of 1 M dithiothreitol (DTT). The transformation mix was first added to pelleted *C. albicans* cells, followed by the addition of the linearized plasmid, and the reactions were left to incubate at 30°C for 1 h. When building the CRISPR libraries, around 20 transformations of the corresponding pooled plasmid library were performed in parallel to increase the number of transformants and achieve higher coverage of all of the plasmid species. The solutions were then heat shocked at 42°C for 50 min in a water bath. Pelleted heat-shocked cells were then washed with 1 mL of fresh YPD three times, then transferred into 9 mL of YPD (for a total of 10 mL) and incubated for ∼3 h at 30°C (∼250 rpm) to allow for expression of the NATr construct. Transformed cells were then pelleted, resuspended in ∼200 µL of YPD, and plated on YPD + NAT (250 µg/mL). Plates were grown at 30°C for 2 days, and a few transformants were then patched onto plates of YPD + NAT (250 µg/mL) and grown at 30°C for an additional day. The patches were PCR tested for the presence of the corresponding sgRNA again, as described earlier. Transformants were also tested for the correct integration into *NEUT5L* by using one primer that binds to the *NEUT5L* region and one that binds to the plasmid.

During CRISPR library construction, all of the heat-shocked and washed cells were mixed together into a single flask for the ∼3 h outgrowth incubation. Afterwards, the pelleted mixture was resuspended in ∼15 mL of YPD, and 100 µL was plated onto YPD + NAT and grown static at 30°C for two days to allow for a rough estimation of transformation efficiency. The remaining volume was inoculated into 1 L of cooled semi-solid YPD media prepared with 3.5 g of SeaPrep^TM^ Agarose from Lonza (cat. 50302) and 250 µg/mL NAT to help minimize jackpotting events and competition between transformants^33,63^. The media was stirred and aliquoted out into three 3 L fernbach flasks as we have noticed *C. albicans* transformants grow poorly beneath a relatively shallow layer of media, set on ice for ∼1 h, and then transferred to incubate static at 30°C for 2-3 days until visible colonies appeared. Following this, the cultures were mixed together into one flask, pelleted, washed with YPD twice to remove residual agarose, and resuspended in ∼10 mL of YPD. The final resuspended libraries were then mixed 1:1 with 40% glycerol and frozen in 2 mL aliquots at -80°C.

### Competitive growth CRISPR-dCas screening

All competitive pooled growth screens were performed in biological quadruplicate to maximize the number of significant strains from the experiment, and to therefore avoid missing key genes underlying the ChrR-mediated drug tolerance^66^. The CRISPRa and CRISPRi screens were both performed identically. Genome copy number was calculated at each step for the given inoculum to help achieve proper representation of all of the mutant types in the library. Four 15 mL YPD + 250 µg/mL NAT cultures of the corresponding thawed CRISPR-dCas library were prepared to a starting OD_600_ of 0.05 and left to grow at 37°C (∼250 rpm) overnight. The following morning, the four cultures were each diluted to an OD_600_ of 0.05 into one tube of 15 mL YPD, one tube of 15 mL YPD + 1 µg/mL FLZ, and one tube of 15 mL YPD + 64 µg/mL FLZ, for a total of 12 cultures. We passaged a blank tube of YPD in parallel to ensure there was no contamination in the media. All cultures were left to grow at 37°C (∼250 rpm) for exactly 12 h. Following this period, the 12 cultures were diluted to an OD_600_ of 0.05 in a fresh 15 mL aliquot of YPD with the same corresponding concentration of FLZ and left to grow at 37°C (∼250 rpm) for a second period of 12 h. This step was repeated for a third 12 h period. After the cultures were grown effectively for 36 h, they were pelleted, resuspended in the remaining ∼2 mL of media after discarding the majority of the supernatant, and frozen at -80°C. The OD_600_ values of the cultures throughout the experiment can be seen in **TABLE S1**. Total generation numbers for all cultures were matched between conditions (within ∼3 generations), thus minimizing potential influences of drift and time-dependent changes in sgRNA abundance.

### gDNA extractions

Pelleted cultures were frozen at -80°C, thawed, and resuspended in 600 μL of homemade sorbitol buffer, which was prepared in nuclease-free water with 1 M sorbitol, 0.1% 2-mercaptoethanol, 0.01 M EDTA, and ∼200 units of zymolase from BioShop (cat. ZYM001.1) per reaction. The resuspended pellets were then left at 35°C (static) for 2 h to generate spheroplasts. Following this, the PureLink™ Genomic DNA Mini Kit from Thermo Fisher Scientific (cat. K182001) was used broadly as per the manufacturer’s instructions. Briefly, spheroplasts were pelleted at 4000 x *g* for 10 min, resuspended in 180 μL of PureLink™ Genomic Digestion buffer and 20 μL of Proteinase K, and incubated at 55°C (static) for 45 min. 20 μL of RNase A (20 mg/mL) was then added to the lysates before being incubated at room temperature for a few minutes. Finally, 200 μL of PureLink™ Genomic Lysis/Binding Buffer and 200 μL of 100% EtOH was added to each sample. Samples were then transferred to a PureLink™ Spin Column and washed/isolated as per the manufacturer’s instructions.

### NGS library prep and sequencing

NGS library preparation was performed in two successive PCR reactions. First, primers with appended homology to the Illumina sequencing adaptors were used to amplify the sgRNA barcode region in the plasmids. Briefly, 5 μL of sample DNA (25 ng for plasmid libraries and 300 ng for fungal libraries) was mixed with 25 μL of NEBNext® High-Fidelity 2X PCR Master Mix from NEB (cat. M0541L), 2.5 μL of each primer (see **TABLE S4**) at 10 μM, and nuclease-free water to 50 μL. Each reaction was set up in duplicate per sample, and PCR-amplified in a thermocycler using the following conditions: 1 cycle of 98°C for 30 s, 25 cycles of 98°C for 10 s followed by 61°C for 30 s and 72°C for 30 s, then a final single cycle of 72°C for 2 min. PCR products were then cleaned up using the NucleoMag kit for clean up and size selection of NGS library prep reactions from Macherey-Nagel (via D-MARK Biosciences) (cat. 744970.5) as per the manufacturer’s instructions. Briefly, the duplicate samples were each pooled and mixed with nucleomag beads in a 1:1.8 ratio of DNA to beads. After incubation at room temperature for 5 min, the beads were separated with a magnet, and the supernatant was discarded. The beads were then washed on a magnet twice with 80% ethanol, and left to dry for ∼15 min until all the leftover ethanol evaporated. After this, the beads were resuspended in 50 μL of nuclease-free water, incubated at room temperature for 5 min, separated with a magnet, and the supernatant containing the cleaned-up product was moved to a new tube. After this, samples were quantified via NanoDrop using an Infinite 200 PRO microplate reader (Tecan) and run on a gel to verify the presence of the correct amplicon. Next, the complete Illumina index primers were appended to each sample by mixing 5 μL of cleaned-up product with 25 μL of the NEBNext® High-Fidelity 2X PCR Master Mix, 5 μL of an Illumina index primer pair at 10 μM, and nuclease-free water to 50 μL. Each reaction was set up in triplicate for each sample, and PCR-amplified in a thermocycler using the following conditions: 1 cycle of 98°C for 30 s, 8 cycles of 98°C for 10 s followed by 55°C for 30 s and 72°C for 30 s, then a final single cycle of 72°C for 2 min. The PCR products were then cleaned up in the same way as after the first reaction, except with a 1:1 ratio of DNA to beads. The samples were eluted again in 50μL of nuclease-free water and run on a gel to verify the correctly-sized libraries and absence of other amplicons.

The final libraries were tested via Nanodrop and Qubit before being diluted to 5 ng/μL, pooled, and sent for sequencing. Libraries were sequenced with the Illumina NovaSeq X platform by SeqCenter in Pittsburgh, PA, except for sequencing of the CRISPRa and CRISPRi plasmid libraries which was done with the Illumina MiSeq and Illumina NovaSeq 6000 platforms by the Advanced Analysis Centre at the University of Guelph. The raw sequencing files have been deposited to the SRA (BioProject ID: PRJNA1471677).

### CRISPR-dCas screening analysis

Our pipeline for analyzing the CRISPR-dCas screening data is available on our GitHub: https://github.com/TheShapiroLab/ChromosomeR, and is similar to the pipeline we have previously described^33^, also available on our Github: https://github.com/TheShapiroLab/Calbicans_Pooled_CRISPRi_2025. Raw sequencing reads were first trimmed, assembled, and aggregated using VSEARCH^67^ and PEAR^68^. Processed reads were assigned to sgRNAs in the library by perfect-match detection of the sgRNA sequence along with 5-nucleotide flanking anchor sequences. To get a sense of how many sgRNAs were captured in our fungal libraries for quality check purposes, sgRNAs needed to be counted at least 50 times in all 4 replicates to be considered as captured in the library (**FIGURE S3**), similarly to our previous work^33^.

During CRISPR screening analysis, sgRNA frequencies were then calculated by first adding a pseudocount of 1 to all counts prior to normalization (to avoid undefined frequencies for zero-count sgRNAs). The list of sgRNAs were filtered by requiring each sgRNA to have a frequency above or equal to the 5th percentile of pseudocount-adjusted frequencies in that sample plus the frequency equivalent of a single count, in at least 3 of 4 replicates in every condition (TP0 YPD, TP3 YPD, TP3 low fluconazole, TP3 high fluconazole). The additional single-count was necessary because many sgRNAs had zero counts at TP3, meaning the 5th percentile would be at the pseudocount floor, such that a percentile-only cutoff would remove nothing. In all cases for the four TP0 replicates, this threshold exceeded the 50 counts we previously used above to define whether an sgRNA was simply present in the library (mean = 180.4 counts across the four TP0 replicates, range 55-321), thus providing increased stringency during screening analysis. We filtered these very low-abundance sgRNAs even in the TP3 samples as we could not confidently verify whether the corresponding depletion was indeed due to a true reduction of fitness and not a result of the population bottleneck during the screen, especially as some of the non-targeting sgRNAs also dropped out by the end of the screen. However, the threshold for the TP3 samples represented a very low absolute number of counts (only 2 counts required in all replicates). Therefore, cases of dramatic depletion were still captured, but only in situations where an sgRNA was still observed in 3/4 of the corresponding TP3 replicates. For each passing sgRNA, log2 FCs were calculated by applying a log2 transformation of the ratio of mean TP3 to mean TP0 replicate frequencies (for the passing replicates only). Significance was assessed using a two-sample t-test (TP3 vs. TP0 replicates) with Benjamini-Hochberg FDR correction applied per condition. An sgRNA was called significant if it had both an FDR < 0.05 and a log2 FC outside the 95% reference interval of the non-targeting sgRNA control distribution (mean ± 1.96 SD). All of the corresponding adjusted p-values and log2 FC values for each sgRNA in each condition are listed in **TABLE S1**.

### Individual and multiplexed CRISPR-dCas plasmid cloning

Following CRISPR screening, individual CRISPR-dCas9 and CRISPR-dCas12a strains were constructed. For single-gene targeting strains, sgRNAs were ordered as a forward and reverse-complement single-stranded DNA oligo pair from Integrated DNA Technologies (IDT), and resuspended to 100 µM in Nuclease Free Duplex Buffer from IDT (cat. 11-05-01-12). For duplexing, oligos were heated at 94°C for 1 min, mixed 1:1 with their counterpart oligo, and heated again at 94°C for 2 min. The duplexed product was then cloned into the corresponding plasmid(s) via Golden Gate with the following mix: ∼1000 ng of plasmid, 1 µL of duplexed oligo, 2 µL of 10X rCutSmart buffer, 2 µL of ATP, 1 µL of SapI, 1 µL of T4 DNA ligase, and nuclease-free water to 20 µL. Reactions were incubated in a thermocycler at 37°C for 2 min followed by 16°C for 5 min for 99 cycles, then 65°C for 15 min, and 80°C for 15 min. After this, an additional 1 µL of SapI was added to each reaction and incubated at 37°C for 1 h to remove any plasmids not containing the properly cloned crRNA. For multiple-gene targeting strains, crRNA arrays with homology to the receiving vector(s) were ordered as GenTitan Gene Fragments from GenScript.

For multiplexed crRNA arrays, insert arrays were designed with ∼20 bp of homology to the receiving vector and ordered as GenTitan Gene Fragments from GenScript. They were then resuspended in ∼30 µL of nuclease-free water and quantified via NanoDrop using an Infinite 200 PRO microplate reader (Tecan). Plasmids were treated with both NaeI (cat. R0190S) and PmlI (cat. R0532S) for the constitutively expressing HyperdCas12a systems, or NaeI alone for the inducible CRISPRi-HyperdCas12a system, involving ∼1000 ng of vector, 1 µL of corresponding enzyme, 5 µL of 10X rCutSmart Buffer, and nuclease-free water to 50 µL. All digestions were performed at 37°C for 12 h followed by a 20 min 65°C heat-inactivation step. Digests were then cleaned up with a DNA Clean & Concentrator-5 kit from Zymo Research (cat. D4013). Following this, 100 ng of cleaned-up vector was mixed with the insert CRISPR array sequence at a 3:1 ratio, along with 10 µL of NEBuilder® HiFi DNA Assembly Master Mix (cat. cat. E2621L), and nuclease-free water to 20 µL. Control reactions set up in the same way but with nuclease-free water instead of an insert were also established in each case to allow for the proportion of false-positive transformants to be determined. When cloning the CRISPR array insert that was designed for the inducible CRISPRi-HyperdCas12a system (pRS1244) into the constitutive CRISPRi-HyperdCas12a system (pRS1074), ie. gRS151, 100 ng of the insert was first treated with PmlI (0.5 µL PmlI, 2.5 µL rCutsmart Buffer, 100 ng of DNA, to 25 µL with nuclease-free water) for 15 min at 37°C, followed by a 20 min 65°C heat inactivation step. In all cases, reactions were then finally incubated at 50°C for 30 min.

Plasmids were transformed into 5-alpha Competent *Escherichia coli* cells from NEB (cat. C2987H), plated onto LB media containing AMP (100 µg/mL) and NAT (250 µg/mL), and left to grow at 30°C for one day. Individual transformants were isolated and patched onto LB + AMP + NAT agar plates and grown at 30°C static for an additional day. To verify correct integration of the sgRNA or individual crRNA sequence into the corresponding plasmid, transformant patches were PCR tested for the presence of the corresponding sequence, where one primer in the PCR is one of the single-stranded sgRNA or crRNA oligos, therefore allowing for a direct test of the presence of the gRNA sequence in the cell. For multiplexed CRISPR-dCas12a plasmids, the whole plasmid was sequenced via Oxford Nanopore PromethION 24 by the Advanced Analysis Centre at the University of Guelph.

For the CRISPRa screen follow-up experiments, we cloned one of the non-targeting sgRNAs that exhibited no significant change in growth in any of the CRISPR-dCas screens into the CRISPRa vector, and used the resulting strain as the control. We observed that the empty vector was significantly enriched in the CRISPR screening experiments, as we have observed similarly before^33^, and therefore would not be a suitable control strain. To generate the individual gene mutant fungal strains, the presence of the gRNA sequence in the genome was tested in the same way as when the corresponding plasmid was generated. For cells transformed with a plasmid containing multiple cRNAs in an array, a large region spanning the entire crRNA array was amplified and sent for Sanger sequencing to verify that the gene fragment was correctly integrated into the plasmid. In all cases, correct integration of the plasmid into the *NEUT5L* region was additionally PCR verified.

### Digital droplet PCR and whole genome sequencing

To confirm that the CRISPRi ChrR-trisomic strains retained the ChrR trisomy (and had no other changes in ploidy) following transformation, we validated the strains had retained an extra copy of ChrR and had no other changes in ploidy via both digital droplet PCR (ddPCR) and WGS. For ddPCR, two primer sets amplifying opposite ends of ChrR (*CR_00980C_A* and *CR_09460C_A*) were used along with a primer set targeting *PMA1* (*C3_00720W_A*) as a control (**TABLE S4**). ddPCR reactions were performed by the Advanced Analysis Centre at the University of Guelph. Briefly, the QX200™ ddPCR™ EvaGreen® Supermix from BioRad (cat. 1864033) was used before droplet generation was performed on a Biorad Automated Droplet Generator using DG8 cartridges and QX200 Droplet Generation Oil for EvaGreen, cycled on a C1000 Touch Thermal Cycler. Droplets were analyzed on a QX200 Droplet Reader. Prior to running the samples, each was diluted by 500-fold and 2 μL was loaded. ddPCR results are available in **TABLE S5**.

Illumina Whole Genome Sequencing (650 Mbp) was performed with the CRISPRi ChrR-trisomic strains (and the euploid CRISPRi control) by SeqCenter in Pittsburgh, PA, on an Illumina NovaSeq X Plus sequencer in one or more multiplexed shared-flow-cell runs, producing 2×151 bp paired-end reads. Raw reads have been deposited to the SRA (BioProject ID: PRJNA1471677). To visualize chromosomal copy number, the Yeast Analysis Mapping Pipeline (YMAP) was used^44^. The raw .fastq files for the strains were uploaded with the ploidy set to 2, paired-end short reads selected, *Candida albicans* SC5314 (CGD, A21-s02-m09-r10) set as the reference genome, and with both GC-content bias and chromosome-end bias options selected. All of the YMAP plots are available in **FIGURE 4B**.

### Minimum inhibitory concentration assays and IC50/supra-MIC calculations

The supra-MIC calculation is a popular method for accurately quantifying antifungal drug tolerance from broth microdilution assays^9,20,25,32^. MIC assays were performed in 96-well flat-bottomed plates. First, 200 μL of the given drug stock was diluted in YPD to twice the highest concentration desired, and added to each well in the first column of the plate. The drug was then serially diluted 1:2 in 100 μL of YPD over the subsequent columns until the second-to-last column. Overnight cultures of *C. albicans* grown in YPD at 30°C (∼250 rpm) were then diluted to an OD_600_ of 0.1 in 1 mL of YPD, and 100 μL of each was then further diluted by 60-fold into YPD. Then, 100 μL of each final diluted culture was mixed into one row each of the 96-well plate, for a final OD_600_ of ∼0.000833. Each plate contained one row of a corresponding *C. albicans* control strain, and a row of blank media as a media contamination control. All strains were tested in triplicate, and plates were read following incubation at 37°C (static) for 24 h and 48 h, unless otherwise stated. OD_600_ values were read using an Infinite 200 PRO microplate reader (Tecan). For the tunicamycin MIC assays, a parallel set of plates with the same starting concentration of DMSO was used to control for the possibility of DMSO-induced growth phenotypes since the concentration of DMSO-solubilized tunicamycin needed in that case was high (**TABLE S3**). Tunicamycin from *Streptomyces* sp. was from Sigma-Aldrich (cat. T7765-5MG).

SMG values were calculated as previously described with some minor modifications^32^. First, we identified the MIC_20_ (the concentration of drug required to inhibit 20% of growth) for each strain replicate by dividing each well by the corresponding no-drug well and checking to see if the resulting value was above 80 after multiplying by 100. We used the MIC_20_ because many highly tolerant strains exhibited high enough SMG levels that a standard MIC_50_ could not be determined. We opted to only consider the wells above the highest MIC_20_ in all strains from the same experiment rather than using their individual MIC_20_ value, as there were many cases of strains growing at a level barely below 80% that of the no-drug well near the MIC_20_, and these values would otherwise artificially inflate the level of tolerance in our CRISPR-dCas strains if kept. Therefore, we averaged the OD_600_ values of all wells above the highest MIC_20_, and divided them by the corresponding no-drug well to produce the SMG value. The statistical significance of the SMG values were calculated using an ordinary one-way ANOVA with Tukey’s multiple comparison test with GraphPad Prism version 10.5.0 for macOS, GraphPad Software, Boston, Massachusetts USA, www.graphpad.com. Raw MIC and SMG data are listed in **TABLE S3** and available on our GitHub: https://github.com/TheShapiroLab/ChromosomeR.

IC_50_ values were calculated by fitting a 4-parameter logistic dose-response curve to the normalized OD values as described previously^32^, via a custom Python script available on our Github: https://github.com/TheShapiroLab/ChromosomeR. FLZ IC_50_ values could not be calculated accurately for the *ACD99* or *PGA13* overexpression strains in the experiment profiling all of the 12 selected follow-up sgRNAs due to their low relative growth in all but the lowest concentration of FLZ, reflecting a very slight decrease in MIC from the other strains, as the other strains grew modestly in the second-to-lowest FLZ concentration. The raw data for both strains can be seen in **TABLE S3**, and do not exhibit any increases in MIC as described in the main text. Potential genetic interactions were calculated using the multiplicative model^40,41^. First, the IC_50_ or SMG values for each strain were normalized to the average value of the corresponding non-targeting control strain, and the product of the individual strains was subtracted from the observed values in the combination strains to calculate ε (deviation from expected). Significance between the expected and observed values was tested for with an unpaired, two-tailed, parametric Welch’s t-test with GraphPad Prism version 10.5.0 for macOS, GraphPad Software, Boston, Massachusetts USA, www.graphpad.com. Genetic interaction analysis results can be seen in **TABLE S3.**

### Reverse transcription quantitative PCR (RT-qPCR)

All RT-qPCR primers used are listed in **TABLE S4**. RT-qPCR was performed with the Luna® Universal One-Step RT-qPCR Kit from NEB (cat. E3005S) using an input of 300-400 ng of RNA (exact amount was kept consistent within each experiment). RNA was first diluted to the corresponding concentration to achieve the desired input, and 5 μL was mixed into a corresponding well on a qPCR 96-well Hard Shell Plate from Applied Biosystems (cat. 4483354) with 15 μL of the master mix prepared as per the Luna® Universal One-Step RT-qPCR Kit with the corresponding RT-qPCR primers. Reactions were performed using a QuantStudio™ 3 Real-Time PCR system from Applied Biosystems. Fold-changes in gene expression were calculated via the comparative C_T_ method ^69^. Briefly, the Cq value of the gene of interest in each RNA sample was subtracted from the housekeeping gene *PMA1* (C3_00720W) to obtain a ΔCt value ^70^. The ΔCt values in the experimental CRISPR-dCas strains were then compared to the corresponding non-targeting control strain to obtain a ΔΔCt value, and finally a fold difference in expression of the target gene(s). The statistical significance when comparing differences in fold-change in target gene expression between strains was calculated using an unpaired, two-tailed, parametric Welch’s t-test with GraphPad Prism version 10.5.0 for macOS, GraphPad Software, Boston, Massachusetts USA, www.graphpad.com. Raw RT-qPCR data are listed in **TABLE S5**.

### Growth curve assays

To perform growth curve assays, overnight cultures of *C. albicans* grown in YPD were first diluted to an OD_600_ of 0.005 in 200 μL of YPD with or without the given stressor in a 96-well plate. Plates were incubated in an Infinite 200 PRO microplate reader (Tecan) at 37°C with orbital shaking for 1000 s in each interval at a 4 mm amplitude. OD_600_ measurements were taken at 20 min intervals over the course of 24 h. The stressors used were 1 M D-Sorbitol from Bioshop (cat. SOR508.1), 1 M Sodium Chloride (NaCl) from Bioshop (cat. SOD001.1), 0.005% Methyl methanesulfonate (MMS) from Sigma-aldrich (cat. 129925-5G), and 3.5 mM Peroxide from FisherScientific (cat. H325-500). For every growth curve assay, all strains were tested in both YPD alone and YPD with the given stressor to control for general stressor-independent fitness defects. Raw growth curve data can be seen in **TABLE S3**, as well as in our GitHub with the corresponding scripts for calculating AUC values: https://github.com/TheShapiroLab/ChromosomeR.

### Assessment of intracellular [3H]-fluconazole accumulation

The assay to measure fluconazole accumulation was performed as published previously^71^. *C. albicans* strains were grown overnight in CSM + 2% glucose at 35°C while shaking. Cells were pelleted and washed three times with PBS and then resuspended in PBS for 3 h for glucose-starvation. Cells were pelleted once more and supernatant discarded, and then 50 µL of concentrated cells were added to 400 µL of PBS without glucose, and 50 µL of freshly diluted 0.77 µM [3H]-fluconazole in a 14 mL round-bottom tube with snap lid. This yielded a final [3H]-fluconazole concentration of 154 nM, which is significantly below the MIC of each strain or isolate being tested. Samples were then incubated with shaking at 30°C for 24 h, after which time the OD_600_ for each sample was recorded. A stop solution (5 mL) (PBS + 20 mM [6 mg/L] unlabeled fluconazole) was added to each sample in the 14 mL tube and the samples were poured over vacuum filtered Whatman glass fiber filters. Cells were washed on the filter by pouring another 5 mL of PBS over the filters and vacuumed until nearly dry. The filters were transferred to a 5 mL scintillation vial along with 3 mL Ecoscint scintillation fluid (Ecoscint XR, National Diagnostics). A Beckman Coulter LS6500 scintillation analyzer (Beckman Coulter, USA) was then used to quantify the radioactivity of each filter as Counts Per Minute (CPM) over the course of 1 min. Experiments were performed in triplicate, and all results were normalized to CPM per 1 × 10^8^ cells. Statistics were performed using an ordinary one-way ANOVA with Šídák’s multiple comparisons test with GraphPad Prism version 10.5.0 for macOS, GraphPad Software, Boston, Massachusetts USA, www.graphpad.com. CPM values are available in **TABLE S3**.

### Rhodamine 6G efflux assays

*C. albicans* strains were analyzed for potential differences in efflux capabilities using the Rhodamine 6G (R6G) assay as previously described^72,73^. This assay measures efflux of R6G from cells into the supernatant over time, and strains that have poor efflux have lower R6G fluorescence than strains with strong efflux capabilities^72^. Strains were initially grown in 50 mL Complete Supplemental Media (CSM) plus 2% glucose to exponential phase (3-5 h) and then were washed with PBS (pH 7) and resuspended in 5 mL PBS + 2 deoxy-D-glucose ((2DG)5 mM). Cell culture OD_600_ was measured and all strain culture ODs were equalized. Cultures were left to incubate at 35°C shaking for 1 h for glucose starvation and temporary efflux inhibition. Cells were then preloaded with R6G at a final concentration of 10 μM (4.8 μg/mL) and allowed to incubate with the fluorescent dye for 30 min. The R6G-preloaded cells were then washed with PBS + 2DG and resuspended in PBS with or without 2% glucose to start the assay (no-glucose data not shown). Efflux of the R6G into the supernatant was measured at time points 0 (immediately after the dye-loaded cells were resuspended in PBS +/- glucose), as well as after 20, 40, and 60 min. This was done by collecting 400 µL of the culture at each time point and transferring to a 1.5 mL microcentrifuge tube, centrifuging at 14000 rpm for 1 min, and transferring 100 µL of supernatant to a black, 96-well plate in duplicate. Finally, the plate was read in a Biotek Synergy H1 Hybrid 96-well plate reader spectrophotometer at 485 nm excitation and 520 nm emission. The data are presented as relative fluorescence units (RFU). All experiments were performed in technical duplicates and repeated to provide biological replicates. The average value across all replicates was plotted for each time point. RFU values are available in **TABLE S3**.

### Sterol extraction and quantification

Sterol extraction and analysis was performed based on the protocol from Arthington-Skaggs et al.^74^ with some modifications. Briefly, each *C. albicans* strain was used to inoculate 10 mL of CSM + 2% glucose. The cultures were incubated at 35°C for 48 h with shaking. The stationary-phase cells were harvested by centrifugation for 3 min and washed once with sterile distilled water. The cell pellet was centrifuged once more for 2 min, and any remaining fluid was discarded by pipetting in order to get a concentrated wet pellet. The net weight of the cell pellet was measured and each strain’s pellet mass was equalized so that ergosterol content could be compared between strains with no additional calculations to account for cell concentration differences. Three milliliters of 25% alcoholic potassium hydroxide solution (25 g of KOH and 36 mL of sterile distilled water, brought to 100 mL with 100% ethanol) was added to each pellet and mixed by vortex for 1 min. Cell suspensions were then transferred to glass tubes with screw top lids and incubated at 85°C for 1 h. Following incubation, tubes were allowed to cool to room temperature. Sterols were then extracted by addition of 1 mL of sterile distilled water and 3 mL of n-heptane, followed by vigorous vortex mixing for 3 min. The tubes were allowed to sit while the layers separated. Two milliliters of the heptane layer containing the sterols was transferred to a clean glass screw-cap tube. Heptane was used as a blank control and 600 µL of each sample was scanned spectrophotometrically between 230 and 300 nm with a Thermo Scientific Evolution 201 UV/Visible Spectrophotometer. The presence of ergosterol and the late sterol intermediate 24(28) dehydroergosterol (DHE) in the extracted samples result in a characteristic four-peaked curve^74^ whereas the absence of detectable ergosterol in extracts and blanks was indicated by a flat line.

Ergosterol content was first normalized across experimental replicates to account for assay variation. Within each replicate, absorbance values for each strain at each wavelength were divided by the corresponding mean across all strains. Normalized A_282_ values, representing the ergosterol peak, were then pooled across the five experimental replicates and expressed as a percentage of the WT NTC control (set to 100%). Statistics were performed using an ordinary one-way ANOVA with Šídák’s multiple comparisons test with GraphPad Prism version 10.5.0 for macOS, GraphPad Software, Boston, Massachusetts USA, www.graphpad.com. Raw spectra data and calculations are listed in **TABLE S3**.

## Supporting information

TABLE S1: Screening Data

TABLE S2: Library Construction

TABLE S3: Growth Profiling Data

TABLE S4: Primer_gRNA_Strain Log

TABLE S5: RT-qPCR and ddPCR Data

## Acknowledgments

The authors would like to thank Dr. Robert Arkowitz, Dr. Iuliana Ene, Dr. Anna Selmecki, and Dr. Petra Vande Zande for helpful discussions and feedback. NCG and MF are supported by NSERC CGS-D awards, LFW was supported by an NSERC PGS-D award, PCD was supported by a Fonds de Recherche du Québec - Santé (FRQS) postdoctoral fellowship (https://doi.org/10.69777/376319), RSS is supported by an NSERC Canada Research Chair. This work was supported by CIHR Project Grant (PJT206047) and NSERC Discovery Grant (RGPIN-2026-04298) to RSS. BioRender images and their availabilities can be found in the respective figure captions.

## Author Contributions

NCG, ACG, and RSS conceptualized the project. NCG, BDE, LFW, MC, and LLH performed the experiments and interpreted the results. NCG, LFW, MF, and PCD contributed to CRISPR-screening analysis design. NCG designed the experiments, analyzed the data, generated the figures, and drafted the manuscript. All authors contributed to revising the manuscript. RSS and TCW supervised the project, provided resources, and acquired funding.

## References

1. Li, R. & Zhu, J. Effects of aneuploidy on cell behaviour and function. Nat. Rev. Mol. Cell Biol. 23, 250–265 (2022).

2. Ben-David, U. & Amon, A. Context is everything: aneuploidy in cancer. Nat. Rev. Genet. 21, 44–62 (2020).

3. Sheltzer, J. M. & Amon, A. The aneuploidy paradox: costs and benefits of an incorrect karyotype. Trends Genet. 27, 446–453 (2011).

4. Vande Zande, P., Zhou, X. & Selmecki, A. The dynamic fungal genome: Polyploidy, aneuploidy and copy number variation in response to stress. Annu. Rev. Microbiol. 77, 341–361 (2023).

5. Gerstein, A. C. & Berman, J. Shift and adapt: the costs and benefits of karyotype variations. Curr. Opin. Microbiol. 26, 130–136 (2015).

6. Case, N. T. et al. Fungal impacts on Earth’s ecosystems. Nature 638, 49–57 (2025).

7. Denning, D. W. Global incidence and mortality of severe fungal disease. Lancet Infect. Dis. 24, e428–e438 (2024).

8. Lee, Y., Robbins, N. & Cowen, L. E. Molecular mechanisms governing antifungal drug resistance. NPJ Antimicrob. Resist. 1, 5 (2023).

9. Berman, J. & Krysan, D. J. Drug resistance and tolerance in fungi. Nat. Rev. Microbiol. 18, 319–331 (2020).

10. Selmecki, A., Gerami-Nejad, M., Paulson, C., Forche, A. & Berman, J. An isochromosome confers drug resistance in vivo by amplification of two genes, ERG11 and TAC1. Mol. Microbiol. 68, 624–641 (2008).

11. Selmecki, A., Forche, A. & Berman, J. Aneuploidy and isochromosome formation in drug-resistant Candida albicans. Science 313, 367–370 (2006).

12. Mackey, A. I. et al. Aneuploidy confers a unique transcriptional and phenotypic profile to Candida albicans. Nat. Commun. 16, 3287 (2025).

13. Tucker, C. et al. Transcriptional regulation on aneuploid chromosomes in divers candida albicans mutants. Sci. Rep. 8, 1630 (2018).

14. Sun, L.-L. et al. Aneuploidy enables cross-tolerance to unrelated antifungal drugs in Candida parapsilosis. Front. Microbiol. 14, 1137083 (2023).

15. Yang, F. et al. Antifungal tolerance and resistance emerge at distinct drug concentrations and rely upon different aneuploid chromosomes. MBio 14, e0022723 (2023).

16. Sun, L.-L. et al. Aneuploidy mediates rapid adaptation to a subinhibitory amount of fluconazole in candida albicans. Microbiol. Spectr. e0301622 (2023).

17. Kukurudz, R. J. et al. Acquisition of cross-azole tolerance and aneuploidy in Candida albicans strains evolved to posaconazole. G3 (Bethesda) 12, (2022).

18. Zheng, L., Xu, Y., Wang, C. & Guo, L. Ketoconazole induces reversible antifungal drug tolerance mediated by trisomy of chromosome R in Candida albicans. Front. Microbiol. 15, 1450557 (2024).

19. Li, X. et al. Trisomy of chromosome R confers resistance to triazoles in Candida albicans. Med. Mycol. 53, 302–309 (2015).

20. Todd, R. T. et al. Antifungal drug concentration impacts the spectrum of adaptive mutations in Candida albicans. Mol. Biol. Evol. 40, msad009 (2023).

21. Jay, A., Jordan, D. F., Gerstein, A. & Landry, C. R. The role of gene copy number variation in antimicrobial resistance in human fungal pathogens. NPJ Antimicrob. Resist. 3, 1 (2025).

22. Ford, C. B. et al. The evolution of drug resistance in clinical isolates of Candida albicans. Elife 4, e00662 (2015).

23. Hirakawa, M. P. et al. Genetic and phenotypic intra-species variation in Candida albicans. Genome Res. 25, 413–425 (2015).

24. Scott, N. E. et al. Aneuploidy, polyploidy and loss of heterozygosity distinguish serial bloodstream isolates of Candida albicans: This article is part of the Candida collection. Microb. Genom. 12, (2026).

25. Rosenberg, A. et al. Antifungal tolerance is a subpopulation effect distinct from resistance and is associated with persistent candidemia. Nat. Commun. 9, 2470 (2018).

26. Lakhani, A. A., Thompson, S. L. & Sheltzer, J. M. Aneuploidy in human cancer: new tools and perspectives. Trends Genet. 39, 968–980 (2023).

27. Gervais, N. C., Hendriks, A. & Shapiro, R. S. Effective strategies for translating CRISPR-dCas systems to diverse microbes. ACS Synth. Biol. 14, 4624–4635 (2025).

28. Chen, G., Bradford, W. D., Seidel, C. W. & Li, R. Hsp90 stress potentiates rapid cellular adaptation through induction of aneuploidy. Nature 482, 246–250 (2012).

29. Linder, R. A., Greco, J. P., Seidl, F., Matsui, T. & Ehrenreich, I. M. The stress-inducible peroxidase TSA2 underlies a conditionally beneficial chromosomal duplication in Saccharomyces cerevisiae. G3 (Bethesda) 7, 3177–3184 (2017).

30. Gervais, N. C. et al. Development and applications of a CRISPR activation system for facile genetic overexpression in Candida albicans. G3 (Bethesda) 13, (2023).

31. Wensing, L., et al. A CRISPR interference platform for efficient genetic repression in Candida albicans. mSphere 4, (2019).

32. Gervais, N. C., Rogers, R. K. J., Robin, M. R. & Shapiro, R. S. HyperdCas12a-based multiplexed genetic regulation in Candida albicans. Nucleic Acids Res. 53, gkaf1402 (2025).

33. Wensing, L. F. et al. Pooled CRISPRi screening reveals fungal-specific drug target candidates. Nat. Microbiol. 1–13 (2026).

34. Wensing, L. et al. Pooled CRISPRi and CRISPRa strain library build for Candida albicans v1. (2026) doi:10.17504/protocols.io.6qpvrb16plmk/v1.

35. Szklarczyk, D. et al. The STRING database in 2023: protein-protein association networks and functional enrichment analyses for any sequenced genome of interest. Nucleic Acids Res. 51, D638–D646 (2023).

36. Lew-Smith, J., Binkley, J. & Sherlock, G. The Candida Genome Database: annotation and visualization updates. Genetics 229, iyaf001 (2025).

37. Chen, H., Zhou, X., Ren, B. & Cheng, L. The regulation of hyphae growth in Candida albicans. Virulence 11, 337–348 (2020).

38. Puerner, C., Serrano, A., Wakade, R. S., Bassilana, M. & Arkowitz, R. A. A myosin light chain is critical for fungal growth robustness in Candida albicans. MBio 12, e0252821 (2021).

39. Silva, P. M., Puerner, C., Seminara, A., Bassilana, M. & Arkowitz, R. A. Secretory vesicle clustering in fungal filamentous cells does not require directional growth. Cell Rep. 28, 2231–2245.e5 (2019).

40. Collins, S. R., Schuldiner, M., Krogan, N. J. & Weissman, J. S. A strategy for extracting and analyzing large-scale quantitative epistatic interaction data. Genome Biol. 7, R63 (2006).

41. Halder, V., McDonnell, B., Uthayakumar, D., Usher, J. & Shapiro, R. S. Genetic interaction analysis in microbial pathogens: unravelling networks of pathogenesis, antimicrobial susceptibility and host interactions. FEMS Microbiol. Rev. 45, fuaa055 (2021).

42. Bouchonville, K., Forche, A., Tang, K. E. S., Selmecki, A. & Berman, J. Aneuploid chromosomes are highly unstable during DNA transformation of Candida albicans. Eukaryot. Cell 8, 1554–1566 (2009).

43. Pfaller, M. A. & Diekema, D. J. Progress in antifungal susceptibility testing of Candida spp. by use of Clinical and Laboratory Standards Institute broth microdilution methods, 2010 to 2012. J. Clin. Microbiol. 50, 2846–2856 (2012).

44. Abbey, D. A. et al. YMAP: a pipeline for visualization of copy number variation and loss of heterozygosity in eukaryotic pathogens. Genome Med. 6, 100 (2014).

45. Basso, L. R., Jr, Gast, C. E., Mao, Y. & Wong, B. Fluconazole transport into Candida albicans secretory vesicles by the membrane proteins Cdr1p, Cdr2p, and Mdr1p. Eukaryot. Cell 9, 960–970 (2010).

46. Calabrese, D., Bille, J. & Sanglard, D. A novel multidrug efflux transporter gene of the major facilitator superfamily from Candida albicans (FLU1) conferring resistance to fluconazole. Microbiology 146 ( Pt 11), 2743–2754 (2000).

47. Marchetti, O. et al. Fungicidal synergism of fluconazole and cyclosporine in Candida albicans is not dependent on multidrug efflux transporters encoded by the CDR1, CDR2, CaMDR1, and FLU1 genes. Antimicrob. Agents Chemother. 47, 1565–1570 (2003).

48. Khandelwal, N. K. et al. Vacuolar sequestration of azoles, a novel strategy of azole antifungal resistance conserved across pathogenic and nonpathogenic yeast. Antimicrob. Agents Chemother. 63, e01347–18 (2019).

49. Baumann, N. A. et al. Transport of newly synthesized sterol to the sterol-enriched plasma membrane occurs via nonvesicular equilibration. Biochemistry 44, 5816–5826 (2005).

50. Klemm, R. W. et al. Segregation of sphingolipids and sterols during formation of secretory vesicles at the trans-Golgi network. J Cell Biol 185, 601–612 (2009).

51. Delic, M. et al. The secretory pathway: exploring yeast diversity. FEMS Microbiol. Rev. 37, 872–914 (2013).

52. Bédard, C. et al. FungAMR: a comprehensive database for investigating fungal mutations associated with antimicrobial resistance. Nat. Microbiol. 10, 2338–2352 (2025).

53. Geng, J., Nair, U., Yasumura-Yorimitsu, K. & Klionsky, D. J. Post-Golgi Sec proteins are required for autophagy in Saccharomyces cerevisiae. Mol. Biol. Cell 21, 2257–2269 (2010).

54. Feng, Y. et al. Body temperature drives azole tolerance in Candida albicans by hindering the autophagic degradation of Erg11. Microbiology (2025).

55. Van Genechten, W. et al. Vacuolar iron export alters the synergy between doxycycline and fluconazole by affecting cidal ROS levels in Candida albicans. MBio 17, e0041626 (2026).

56. Koller, M. S., Himmelbauer, C., Fink, S., Ravichandran, M. C. & Campbell, C. S. Combinatorial effects of multiple genes contribute to beneficial aneuploidy phenotypes. Genetics (2025).

57. Williams, C. R., Baccarella, A., Parrish, J. Z. & Kim, C. C. Trimming of sequence reads alters RNA-Seq gene expression estimates. BMC Bioinformatics 17, 103 (2016).

58. Dobin, A. et al. STAR: ultrafast universal RNA-seq aligner. Bioinformatics 29, 15–21 (2013).

59. Muzafar, S., Sharma, R. D., Chauhan, N. & Prasad, R. Intron distribution and emerging role of alternative splicing in fungi. FEMS Microbiol. Lett. 368, (2021).

60. Liao, Y., Smyth, G. K. & Shi, W. featureCounts: an efficient general purpose program for assigning sequence reads to genomic features. Bioinformatics 30, 923–930 (2014).

61. Robinson, M. D., McCarthy, D. J. & Smyth, G. K. edgeR: a Bioconductor package for differential expression analysis of digital gene expression data. Bioinformatics 26, 139–140 (2010).

62. Peng, D. & Tarleton, R. EuPaGDT: a web tool tailored to design CRISPR guide RNAs for eukaryotic pathogens. Microb. Genom. 1, e000033 (2015).

63. Momen-Roknabadi, A., Oikonomou, P., Zegans, M. & Tavazoie, S. An inducible CRISPR interference library for genetic interrogation of Saccharomyces cerevisiae biology. *Commun*. Biol. 3, 723 (2020).

64. Horlbeck, M. A. et al. Compact and highly active next-generation libraries for CRISPR-mediated gene repression and activation. Elife 5, (2016).

65. Langmead, B., Trapnell, C., Pop, M. & Salzberg, S. L. Ultrafast and memory-efficient alignment of short DNA sequences to the human genome. Genome Biol. 10, R25 (2009).

66. Robinson, D. G., Chen, W., Storey, J. D. & Gresham, D. Design and analysis of Bar-seq experiments. G3 (Bethesda) 4, 11–18 (2014).

67. Rognes, T., Flouri, T., Nichols, B., Quince, C. & Mahé, F. VSEARCH: a versatile open source tool for metagenomics. PeerJ 4, e2584 (2016).

68. Zhang, J., Kobert, K., Flouri, T. & Stamatakis, A. PEAR: a fast and accurate Illumina Paired-End reAd mergeR. Bioinformatics 30, 614–620 (2014).

69. Schmittgen, T. D. & Livak, K. J. Analyzing real-time PCR data by the comparative C(T) method. Nat. Protoc. 3, 1101–1108 (2008).

70. Nailis, H., Coenye, T., Van Nieuwerburgh, F., Deforce, D. & Nelis, H. J. Development and evaluation of different normalization strategies for gene expression studies in Candida albicans biofilms by real-time PCR. BMC Mol. Biol. 7, 25 (2006).

71. Esquivel, B. D. & White, T. C. Transport across membranes: Techniques for measuring drug import in fungal cells. Methods Mol. Biol. 2658, 215–221 (2023).

72. Esquivel, B. D. & White, T. C. Transport across membranes: Techniques for measuring efflux in fungal cells. Methods Mol. Biol. 2658, 201–213 (2023).

73. Esquivel, B. D. et al. Characterization of the Efflux Capability and Substrate Specificity of Aspergillus fumigatus PDR5-like ABC Transporters Expressed in Saccharomyces cerevisiae. MBio 11, e00338–20 (2020).

74. Arthington-Skaggs, B. A., Warnock, D. W. & Morrison, C. J. Quantitation of Candida albicans ergosterol content improves the correlation between in vitro antifungal susceptibility test results and in vivo outcome after fluconazole treatment in a murine model of invasive candidiasis. Antimicrob. Agents Chemother. 44, 2081–2085 (2000).

