## Supplementary Information for "Chromosome-scale CRISPR screening reveals secretory pathway genes as drivers of aneuploidy-mediated antifungal tolerance"

This PDF file includes:

**Part A: Supplementary Figures**
**Part B: Supplementary Methods**
**Part C: Description of Supplementary Tables**
**Supplementary References**

**Part A: Supplementary Figures**

- Figure S1: ChrR trisomies lead to general growth defects and a predictable shift in global gene expression dominated by increased expression of ChrR genes.
- Figure S2: The individual C. albicans transcriptional responses to fluconazole and ChrR aneuploid stress.
- Figure S3: Distributions of and statistics for the plasmid and fungal pooled CRISPR-dCas libraries.
- Figure S4: CRISPR-dCas screening activity correlations by replicates and growth conditions.
- Figure S5: CRISPRi/a-dCas screens in plain nutrient-rich media, relatively low FLZ, and results summary.
- Figure S6: Profiling and validation of selected ChrR genes and their corresponding mutant strains.
- Figure S7: Characterizing the impact of individually repressing or overexpressing SEC4, YPT31, and ACD99 on growth during various stressors.
- Figure S8: Upregulation of SEC4 and YPT31 is sufficient for ChrR aneuploidy-mediated posaconazole tolerance and does not significantly alter azole IC50.
- Figure S9: ACD99 does not likely contribute to ChrR-mediated antifungal tolerance.
- Figure S10: Upregulation of the secretory pathway is necessary for the heightened ChrR-mediated posaconazole tolerance and high levels of tolerance in clinical isolates.
- Figure S11: Characterizing the influence of SEC4 and YPT31 upregulation on growth during ER stress as well as during growth with fluconazole with or without efflux.

**Part B: Supplementary Methods**

- Principal Component Analysis and PERMANOVA
- Multi-Omics Integration and Candidate Gene Selection
- STRING Analysis
- Clamped Homogeneous Electric Field (CHEF) Electrophoresis

**Part C: Description of Supplementary Tables**

**Supplementary References**

### Part A: Supplementary Figures

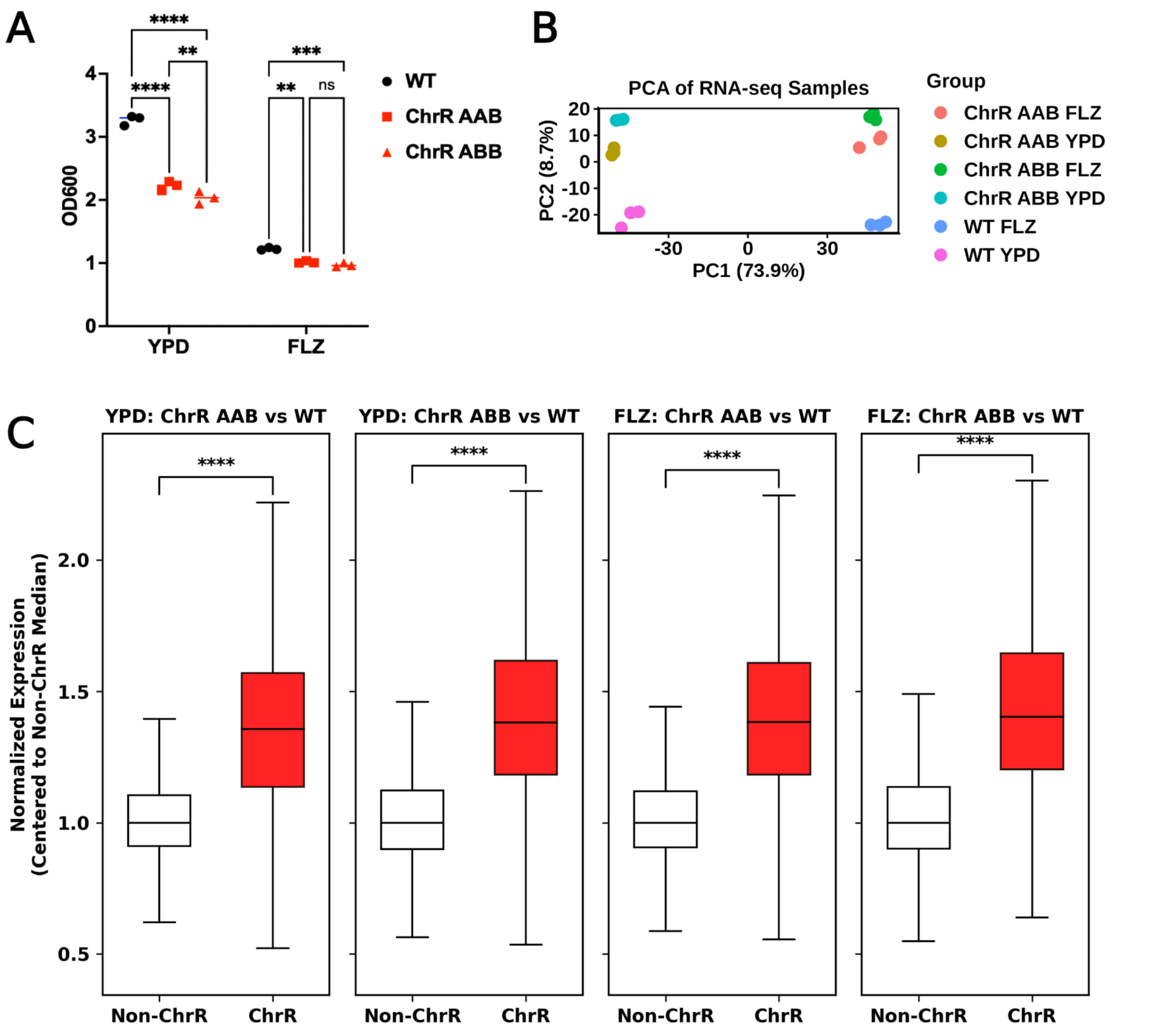


**Figure S1: ChrR trisomies lead to general growth defects and a predictable shift in global gene expression dominated by increased expression of ChrR genes.**

**(A)** $OD_{600}$ values of the euploid WT and the two ChrR-trisomic strains after 6 h of growth at 37°C in either plain nutrient-rich media (YPD) or the same media supplemented with 64 µg/mL of FLZ (starting OD600 was 0.05). Statistics were performed with an ordinary two-way ANOVA and Šidák's multiple comparisons test, $^{**}P < .01$, $^{***}P < 0.001$, $^{****}P < .0001$. **(B)** Principal component analysis (PCA) of the RNA-seq samples by genotype and treatment. PERMANOVA analysis revealed that both drug treatment ($R^2 = 0.73$, $P = 0.001$) and aneuploidy ($R^2 = 0.13$, $P = 0.001$) contributed significantly to global variance, though fluconazole accounted for 5.6-fold more variance in transcription. There was no significant interaction between FLZ and presence of the ChrR trisomy in the PERMANOVA ($R^2 = 0.024$, $P = 0.144$), suggesting that while the presence of the ChrR trisomy uniquely alters the expression of thousands of genes (**FIGURE 1E**), the overall magnitude of the transcriptional response to FLZ is not significantly different from that of the euploid strain. **(C)** The average normalized expression of ChrR genes in ChrR-trisomic strains in both plain nutrient-rich media and drug stress compared to the normalized expression of non-ChrR genes in ChrR-trisomic strains. Statistics were performed using a two-sided Mann-Whitney U test to compare the ChrR and non-ChrR gene expression distributions, *$P < .05$, ****$P < .0001$.

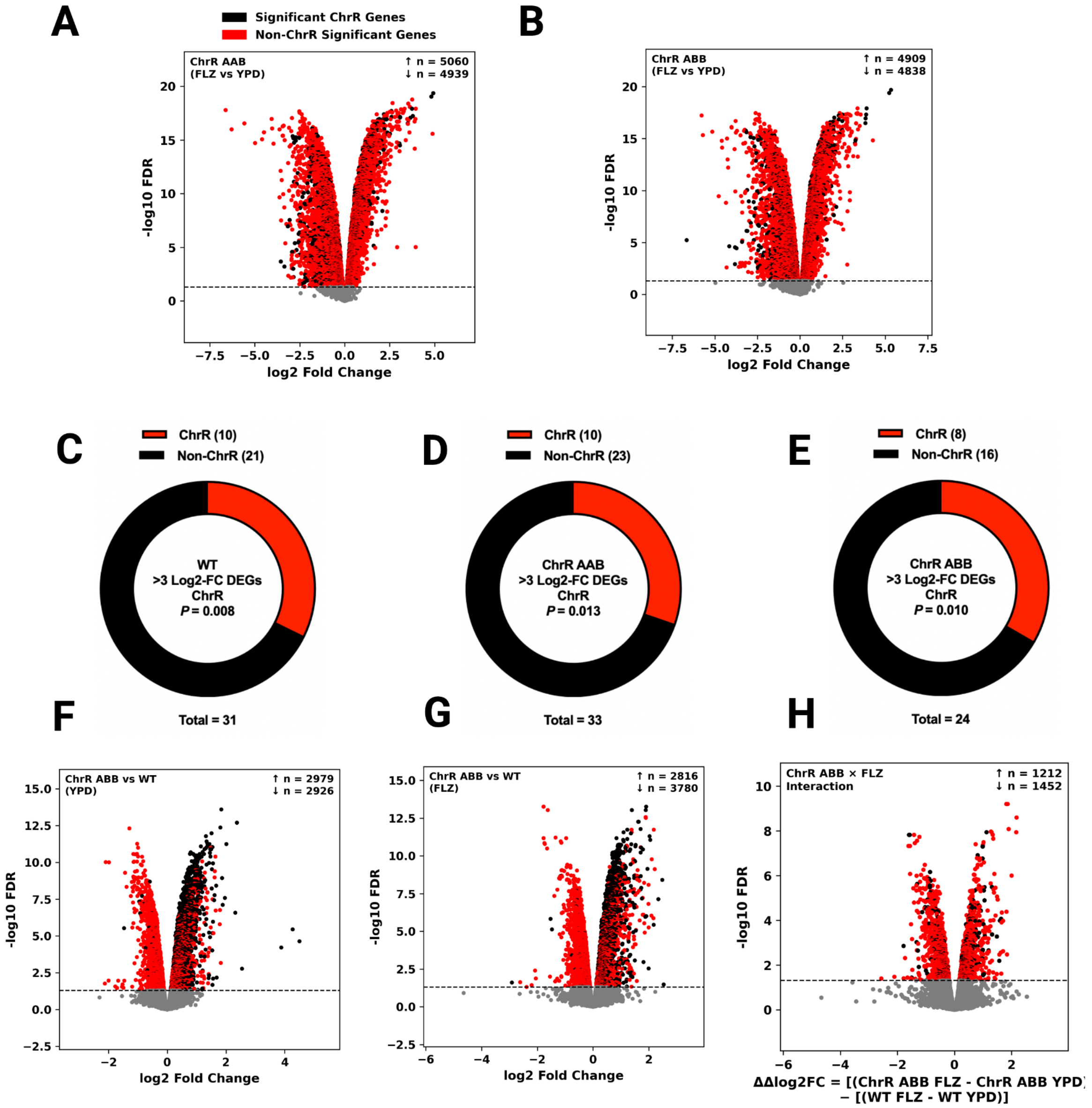


**Figure S2: The individual *C. albicans* transcriptional responses to fluconazole and ChrR aneuploid stress.**

**(A)** Global gene expression changes in ChrR-trisomic strains ChrR AAB and **(B)** ChrR ABB when grown in YPD supplemented with 64 µg/mL FLZ compared to plain YPD. **(C)** The proportion of ChrR genes that were highly upregulated (>3 log2 FC) in the euploid WT, **(D)** ChrR AAB, and **(E)** ChrR ABB, when grown in YPD supplemented with 64 µg/mL FLZ compared to plain YPD. The significance of the proportion of strongly upregulated genes on ChrR was tested using a hypergeometric test (testing the significance of the number of >3 log FC genes on ChrR based on the total number of >3 log FC genes on every chromosome, the number of genes on

ChrR, and the number of genes in the genome). Notably, all three strains shared a >3 log2 FC upregulation of several ChrR genes, including *INO1* (*CR_10100C*), *CR_03580C*, and *HXT5* (*CR_03450W*), suggesting they may play a role in response to a high dose of FLZ (**TABLE S1**). **(F)** Changes in global gene expression in the ChrR-trisomic strain ChrR ABB compared to the euploid WT when both were grown in plain YPD and **(G)** when both were grown in YPD supplemented with 64 µg/mL FLZ. **(H)** The aneuploid-conditioned FLZ response, ie. the interaction between the ChrR trisomy and FLZ stress response in the ChrR-trisomic strain (ChrR ABB) (See Methods).

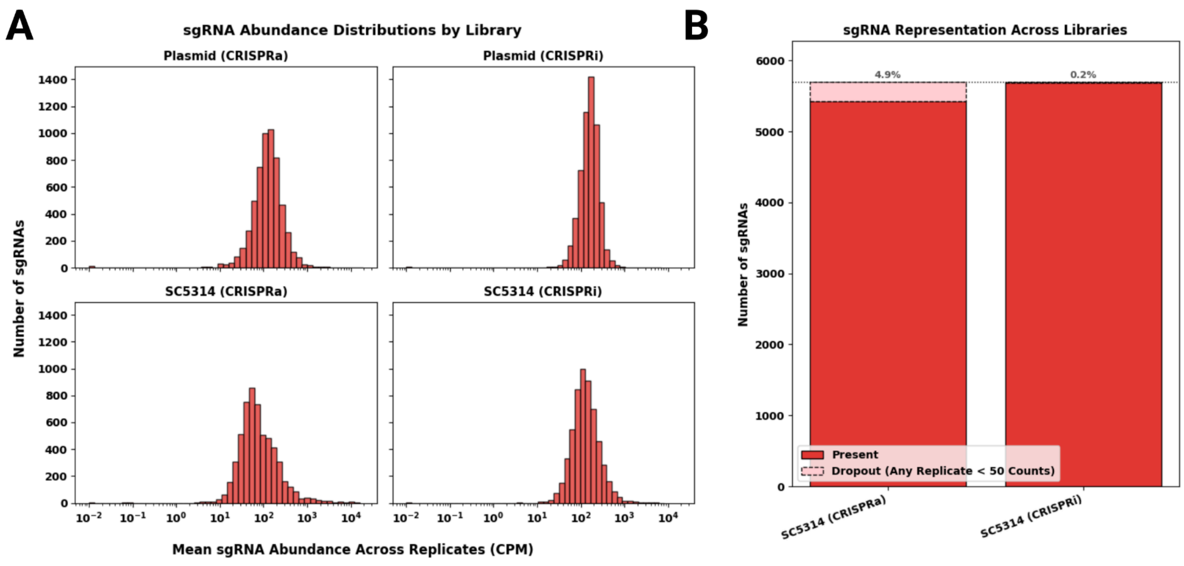


**Figure S3: Distributions of and statistics for the plasmid and fungal pooled CRISPR-dCas libraries.**

**(A)** 5700 sgRNAs were designed to target all genes on ChrR, synthesized as an oligo pool, and cloned into our CRISPRa-dCas9[1] and CRISPRi-dCas9[2] plasmid backbones. A region encompassing the 20bp sgRNA barcode sequence in the corresponding plasmid and fungal (*C. albicans* SC5314) libraries was amplified and sequenced in triplicate (plasmid) or quadruplicate (fungal), and each barcode sequence (flanked by 8bp constant regions) in each replicate sample was then searched for and counted. The resulting absolute counts were used to calculate a count per million (CPM) by dividing the absolute count by the total number of counted sgRNAs and multiplying by 1,000,000. To determine how many sgRNAs were captured in the final fungal libraries, each sgRNA had to be counted at least 50 times in all corresponding replicates, as previously described[3]. **(B)** The total number of sgRNAs present in each library is listed along with the number of sgRNAs that were not detected (%).

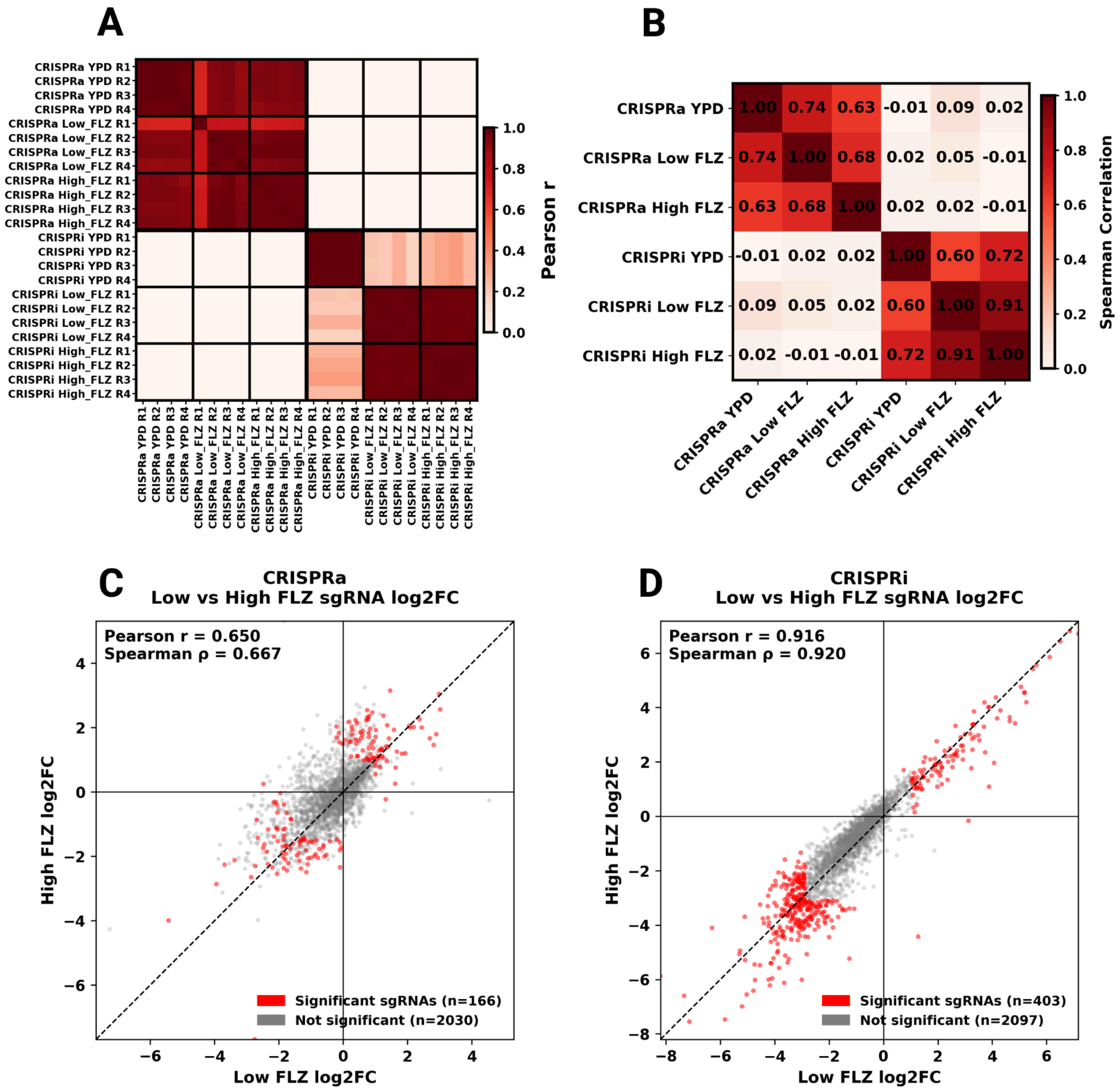


**Figure S4: CRISPR-dCas screening activity correlations by replicates and growth conditions.**

**(A)** Correlation matrix of the CRISPRi/a-dCas screens by replicate. Pearson r was used to assess the correlation between guide frequencies to quantify reproducibility across replicates. **(B)** Correlation matrix of the CRISPRi/a-dCas screens by condition per system. Spearman ρ was used on log2 FC values to quantify the rank-order similarities between screening conditions. **(C)** Correlation plot between the 1 μg/mL and 64 μg/mL FLZ conditions in the CRISPRa screens and the **(D)** CRISPRi screens. Only sgRNAs that were counted (ie. had a corresponding log2 FC and adjusted p-value) in both concentrations of FLZ at the end of the corresponding screen are included. Significant sgRNAs correspond to an sgRNA that had an adjusted p-value of < 0.05 as well as a log2 FC outside of the corresponding 95% confidence interval (see Methods) in at least one of the FLZ concentrations.

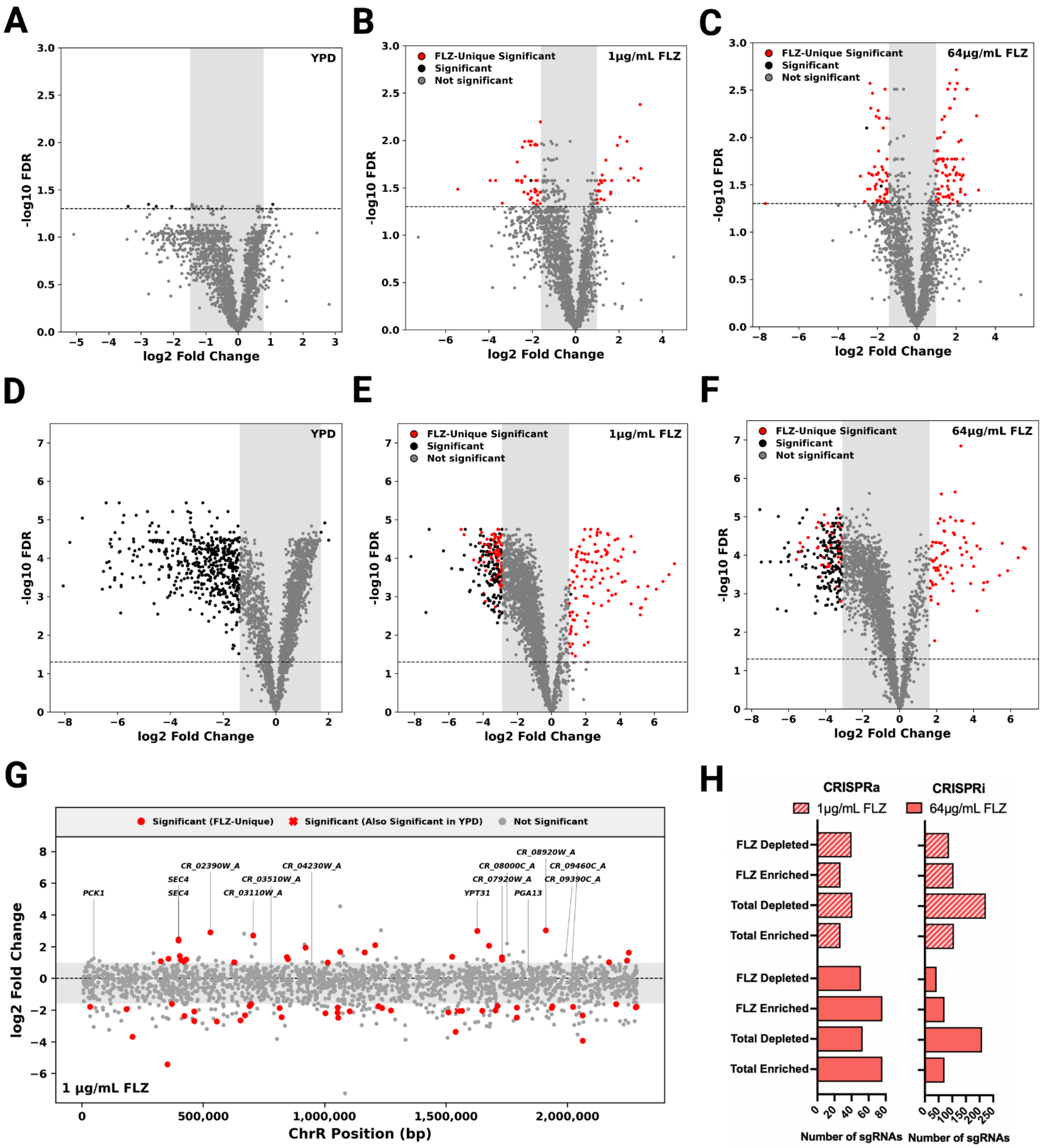


**Figure S5: CRISPRi/a-dCas screens in plain nutrient-rich media, relatively low FLZ, and results summary.**

**(A)** The changes in sgRNA abundance in the CRISPRa screen in YPD alone, **(B)** the CRISPRa screen in 1 µg/mL FLZ, **(C)** the CRISPRa screen in 64 µg/mL FLZ, **(D)** the CRISPRi screen in YPD alone, **(E)** the CRISPRi screen in 1 µg/mL FLZ, and **(F)** the CRISPRi screen in 64 µg/mL FLZ. The grey shaded areas represent 95% confidence intervals based on the log2 FCs of the non-targeting sgRNAs in each pooled library for the given condition. As expected, passaging in YPD alone produced substantially more changes in relative strain abundance in the CRISPRi library than in the CRISPRa library, likely reflecting the widespread fitness consequences of repressing critical or essential genes. In the FLZ screens, the sgRNAs that were uniquely significantly differentially abundant in the FLZ screen (and not significant in the corresponding YPD-alone screen) are emphasized (red). **(G)** The changes in sgRNA abundance in the CRISPRa screen in 1 µg/mL FLZ. Each datapoint represents an sgRNA at its corresponding target position along the length of ChrR. The grey shaded area represents a 95% confidence interval based on the log2 fold-changes of the non-targeting sgRNAs. sgRNAs that were significantly enriched or depleted when the library was grown in plain YPD are marked with “X” since they likely do not represent changes specific to the 1 µg/mL FLZ environment. **(H)** Sum of the number of sgRNAs that were enriched or depleted with both systems in 1 µg/mL FLZ and 64 µg/mL FLZ beyond the corresponding 95% confidence interval set by the non-targeting sgRNAs. Candidate genes proposed to potentially underlie ChrR-mediated drug tolerance vary substantially across studies. *GZF3* (*CR_02850C*) has previously been identified as a potential mediator of antifungal tolerance[4,5], though we did not identify a clear relationship between *GZF3* and antifungal tolerance in our CRISPR-dCas screens (**TABLE S1**). *GZF3* was also very weakly upregulated in FLZ in one of the ChrR trisomic strains, though it was upregulated by ~2.0-fold in the other (**TABLE S1**). The ChrR gene *MNL1* has previously been demonstrated to enhance azole tolerance when either deleted or overexpressed[6]. While we did not observe the latter, one sgRNA targeting *MNL1* was indeed one of the most significantly enriched in both fluconazole conditions (>4-fold log2 FC) in our CRISPRi screens (**TABLE S1**). Therefore, there may be many ChrR genes that can lead to antifungal tolerance that may or may not be directly caused by gene upregulation via ChrR aneuploidy.

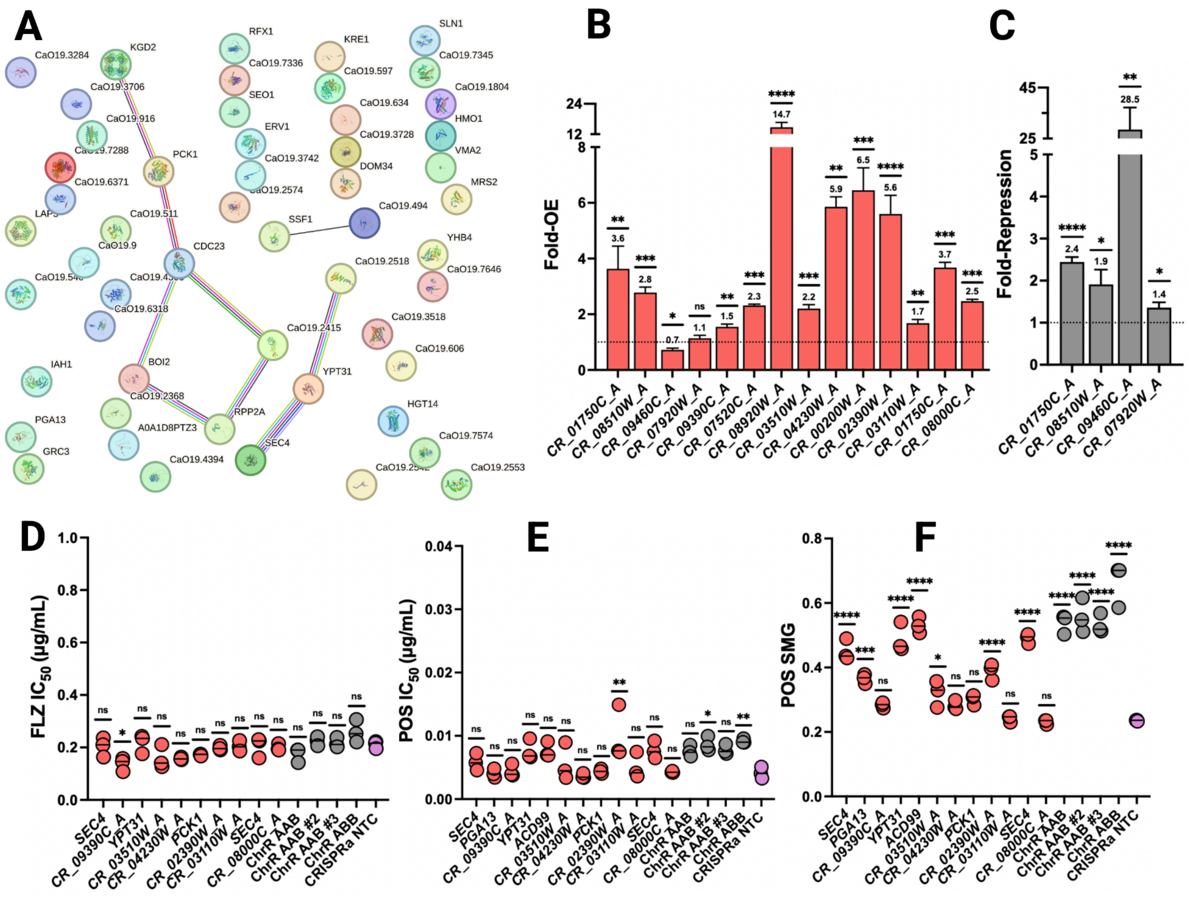


**Figure S6: Profiling and validation of selected ChrR genes and their corresponding mutant strains.**

**(A)** STRING analysis[7] was performed with all of the proteins corresponding to the enriched sgRNAs in the CRISPRa FLZ screens. Circles represent nodes/proteins, and lines represent edges and indicate interactions between proteins. The number of edges between proteins represents different types of interactions between proteins. Our STRING analysis highlighted the strong connection between Sec4 and Ypt31. A second small module centered around *CR_08630W* (*CDC23*) also formed, though the functions of the genes in the corresponding network were diverse and did not represent a single clear pathway. However, *CDC23* shares a role in the anaphase-promoting complex with *CDC28*, the latter of which is required for proper localization of Sec2, which activates Sec4 after being recruited by Ypt31[8,9]. Therefore, this secondary module may also affect secretory-mediated azole tolerance. **(B)** The level of differential expression of the listed target gene in the CRISPRa-dCas9 strains when they were reconstructed with the selected sgRNAs from the screens. Statistics were performed by comparing the dCt values between the given strain and its corresponding non-targeting control strain using a Welch's two-tailed t-test, ns $P > 0.05$, *$P < 0.05$, **$P < 0.01$, ***$P < 0.001$, ****$P < 0.0001$. The two strains that were not overexpressing their target gene were either significantly repressing the target, which we have observed with CRISPRa in the past [10], or could have potentially overexpressed the gene in fluconazole during the screen but not plain YPD if the transcriptional start site (and therefore on-target CRISPR-dCas activity) was dependent on the environmental condition [1], if it did not represent a true false positive. **(C)** The level of repression of the listed target gene in the CRISPRi-dCas9 strains when they were reconstructed with the selected sgRNAs from the CRISPRi screens. Statistics were performed by comparing the dCt values between the given strain and its corresponding non-targeting control strain using a Welch's two-tailed t-test, *$P < 0.05$, **$P < 0.01$, ****$P < 0.0001$. **(D)** The $IC_{50}$ in FLZ and **(E)** the $IC_{50}$ in POS, and **(F)** the SMG (ie. average growth above the $MIC_{20}$ at 48 h) in POS in the individual overexpression (CRISPRa) strains selected from the screens. Statistics were performed using an ordinary one-way ANOVA with Tukey's multiple comparison test, where significance represents the difference between the given strain and the non-targeting control strain (NTC), ns $P > 0.05$, *$P < 0.05$, ***$P < 0.001$, ****$P < 0.0001$. Many of the subtle increases in azole tolerance in FLZ were not significant in POS, possibly because the overall tolerance level among the tested strains was lower in POS than in FLZ.

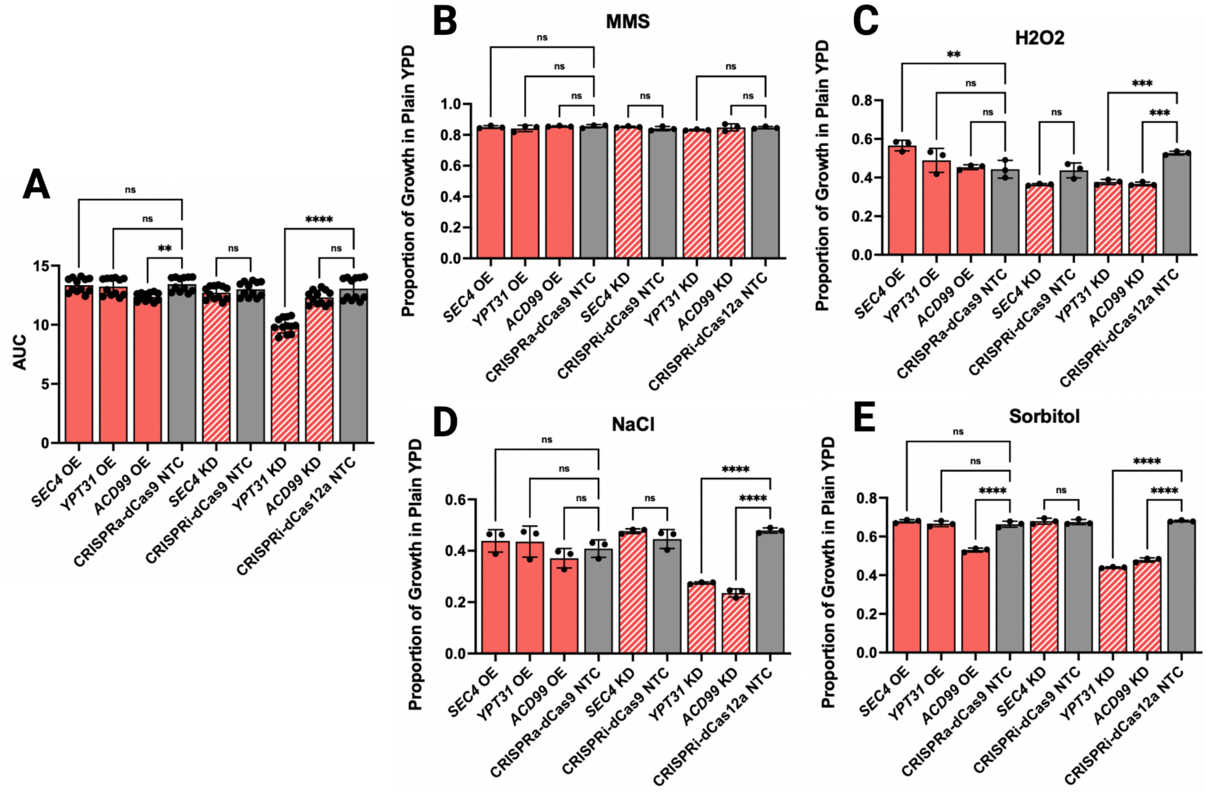


**Figure S7: Characterizing the impact of individually repressing or overexpressing *SEC4*, *YPT31*, and *ACD99* on growth during various stressors.**

**(A)** Area under the curve (AUC) values were calculated and combined from all 4 stressor growth curve assays in plain YPD alone. These assays included the corresponding individual overexpression (OE) or knockdown (KD) strains of the three genes, along with the corresponding non-targeting control (NTC) strains. **(B)** AUC values were calculated from the 4 growth curve assays in the given stressor and compared to the AUC values in plain YPD in the matched experiment. The stressors tested included 0.005% methyl methanesulfonate (MMS), **(C)** 3.5 mM Peroxide (H2O2), **(D)** 1 M Sodium Chloride (NaCl), and **(E)** 1 M D-Sorbitol. Statistics were performed using an ordinary one-way ANOVA with Tukey's multiple comparison test, ns $P > 0.05$, $^{**}P < 0.01$, $^{***}P < 0.001$, $^{****}P < 0.0001$.

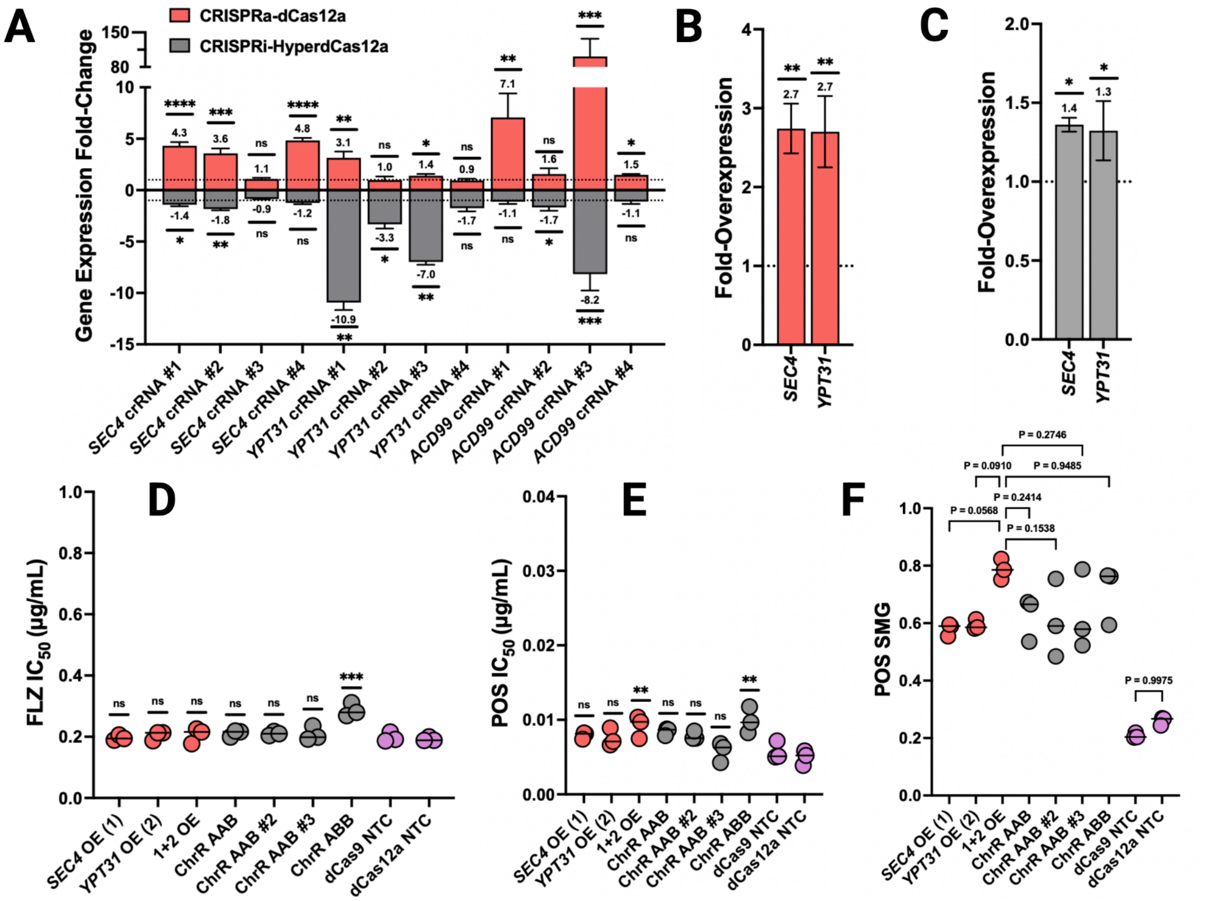


**Figure S8: Upregulation of *SEC4* and *YPT31* is sufficient for ChrR aneuploidy-mediated posaconazole tolerance and does not significantly alter azole $IC_{50}$.**

**(A)** The level of overexpression and repression of the listed target gene when the same set of crRNAs were tested individually in both the constitutively-expressing WT CRISPRa-dCas12a system and the inducible CRISPRi-HyperdCas12a system. Statistics were performed by comparing the dCt values between the given strain and its corresponding non-targeting control strain using a Welch's two-tailed t-test, ns $P > 0.05$, *$P < 0.05$, **$P < 0.01$, ***$P < 0.001$, ****$P < 0.0001$. **(B)** The level of overexpression of *SEC4* and *YPT31* in the multiplexed CRISPRa strain. Statistics were performed by comparing the dCt values between the given strain and the corresponding non-targeting control strain using a Welch's two-tailed t-test, **$P < 0.01$. **(C)** The level of expression of *SEC4* and *YPT31* in the two tested ChrR-trisomic strains (ChrR AAB and ChrR ABB) compared to the euploid WT when grown in FLZ, tested via RNA-seq. Statistical significance reflects the adjusted p-value from the RNA-seq experiments. Minimum and maximum values in error bars represent the values in the two ChrR-trisomic strains. Significance varied per gene in both strains compared to the WT, but was significant in all cases, so for simplicity, *$P < 0.01$. **(D)** The $IC_{50}$ of each strain in FLZ and **(E)** POS. Statistics were performed using an ordinary one-way ANOVA with Tukey's multiple comparison test, ns >

0.05, *P < 0.05, ****P < 0.0001. **(F)** The SMG in posaconazole in the individual and combination overexpression strains. Statistics were performed using an ordinary one-way ANOVA with Tukey's multiple comparison test, where significance represents the difference between the given strain and the non-targeting control strain (NTC), ns *P* > 0.05, ***P* < 0.01. Exact p-values are listed for the POS SMG experiment for improved clarity as some tests approached or barely met the threshold for significance.

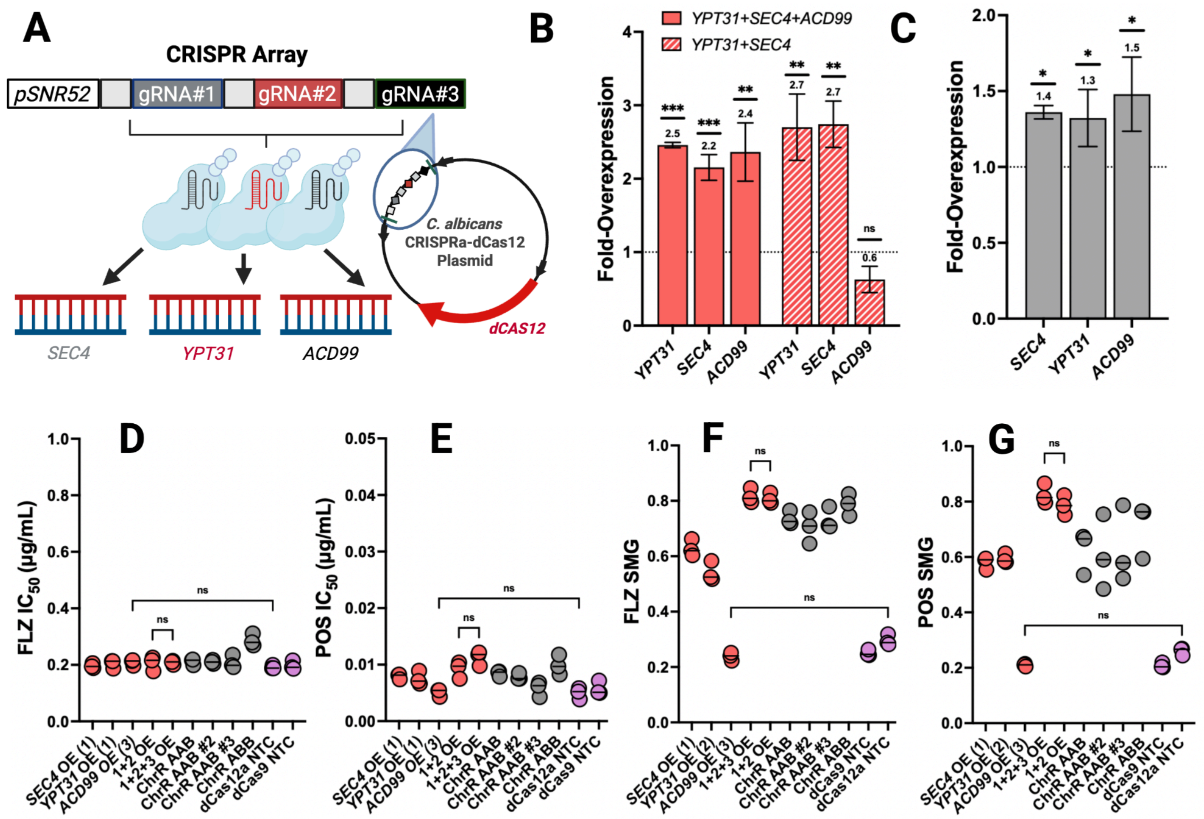


**Figure S9: *ACD99* does not likely contribute to ChrR-mediated antifungal tolerance.**

**(A)** Conceptual schematic of the WT dCas12a-based multiplexed targeting of *SEC4*, *YPT31*, and *ACD99*. **(B)** The level of overexpression of *SEC4, YPT31*, and *ACD99* in the multiplexed CRISPRa strains when all three or just the former two were targeted. Statistics were performed by comparing the dCt values between the given strain and its corresponding non-targeting control strain using a Welch's two-tailed t-test, ns $P > 0.05$, **$P < 0.01$, ***$P < 0.001$. **(C)** The level of expression of *SEC4*, *YPT31*, and *ACD99* in the two tested ChrR-trisomic strains (ChrR AAB and ChrR ABB) compared to the euploid WT when grown in FLZ, tested via RNA-seq. Statistical significance reflects the adjusted p-value from the RNA-seq experiments. Minimum and maximum values in error bars represent the values in the two ChrR-trisomic strains. Significance varied per gene in both strains compared to the WT, but were significant in all cases, so for simplicity, *$P < 0.01$. **(D)** The $IC_{50}$ in both fluconazole and posaconazole **(E)** in the individual and combination overexpression strains. The mildly-overexpressing *ACD99* CRISPRa-dCas12a strain did not lead to an increased level of tolerance, suggesting *ACD99*-mediated antifungal tolerance is dependent on a very high level of overexpression and is thus unlikely to underlie ChrR-mediated antifungal tolerance. Overexpressing *ACD99* also did not further improve the level of tolerance achieved via *SEC4* and *YPT31* upregulation alone.

Statistics were performed using an ordinary one-way ANOVA with Tukey's multiple comparison test, ns $P > 0.05$. **(F)** The SMG for each strain grown in FLZ and **(G)** POS. Statistics were performed using an ordinary one-way ANOVA with Tukey's multiple comparison test, where significance represents the difference between the given strain and the non-targeting control strain (NTC), ns $P > 0.05$.

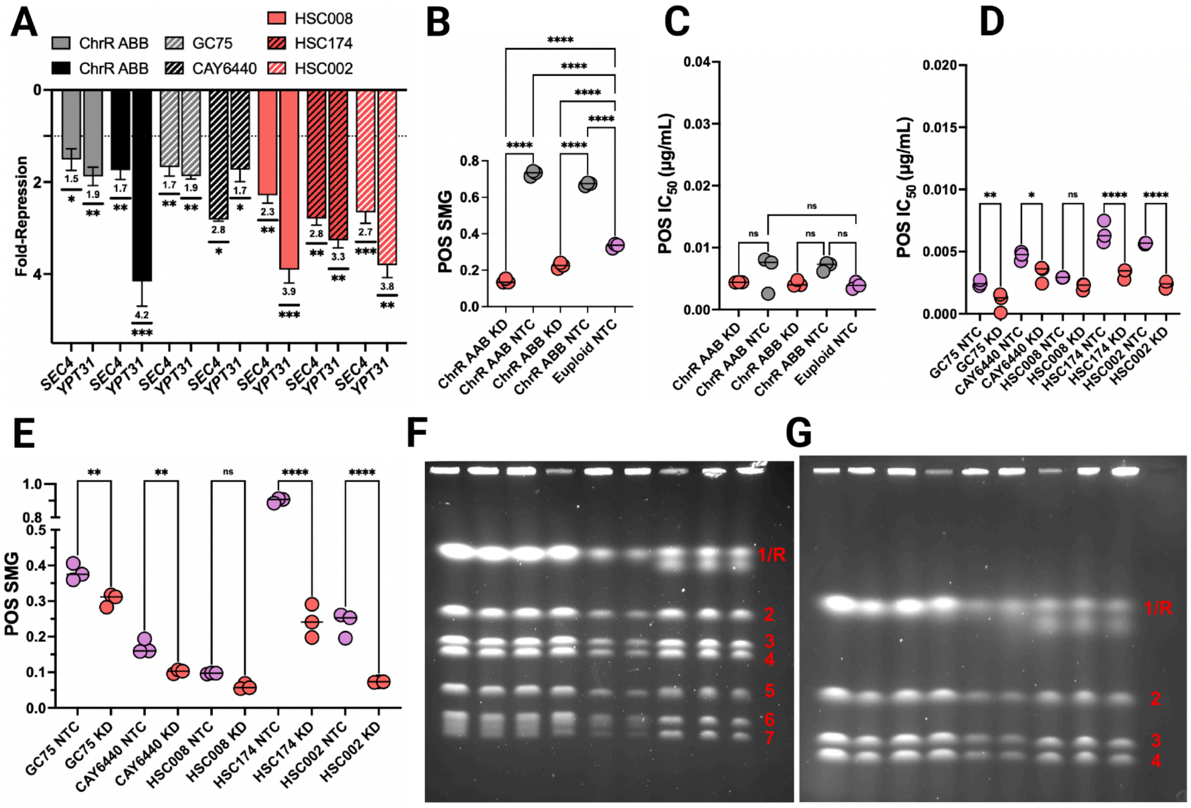


**Figure S10: Upregulation of the secretory pathway is necessary for the heightened ChrR-mediated posaconazole tolerance and high levels of tolerance in clinical isolates.**

**(A)** The level of repression of *SEC4* and *YPT31* in the multiplexed CRISPRi strains. Statistics were performed by comparing the dCt values between the given strain and its corresponding non-targeting control strain using a Welch's two-tailed t-test, $*P < 0.05$, $**P < 0.01$, $***P < 0.001$. **(B)** The SMG and **(C)** $IC_{50}$ of each of the ChrR-trisomic *SEC4*+*YPT31* knockdown (KD) strains and the euploid non-targeting control (NTC) strain in POS. Statistics were performed using an ordinary one-way ANOVA with Šídák's multiple comparisons test, ns $P > 0.05$, $****P < 0.0001$. **(D)** The $IC_{50}$ and **(E)** SMG of each of the clinical isolate *SEC4*+*YPT31* knockdown (KD) strains and the corresponding non-targeting control (NTC) strains in POS. Statistics were performed using an ordinary one-way ANOVA with Šídák's multiple comparisons test, ns $P > 0.05$, $*P < 0.05$, $**P < 0.01$, $****P < 0.0001$. **(F)** *C. albicans* strains SC5314, GC75, and CAY6440 were passaged in duplicate in 64 µg/mL FLZ, and compared to a DMSO-grown control to check for standing ploidy or immediate ploidy differences following brief exposure to high FLZ concentration. Chromosomes were visualized by Clamped Homogeneous Electric Field (CHEF) electrophoresis for a total run time of 99 h. Run time was increased to 144 h **(G)** to attempt resolution of the large chromosomes. Lane order for (F) and (G): SC5314 DMSO, SC5314 FLZ replicate 1, SC5314 FLZ replicate 2, GC75 DMSO, GC75 FLZ replicate 1, GC75 FLZ replicate 2, CAY6440 DMSO, CAY6440 FLZ replicate 1, and CAY6440 FLZ replicate 2. The largest chromosomes (Chr1 and ChrR) were unresolved for SC5314 and GC75, and only partially resolved for CAY6440. Copy number and/or size variation was observed among the strains. For all isolates, FLZ exposure did not affect the CHEF karyotype. Positions of the corresponding chromosomes are labeled 1/R to 7.

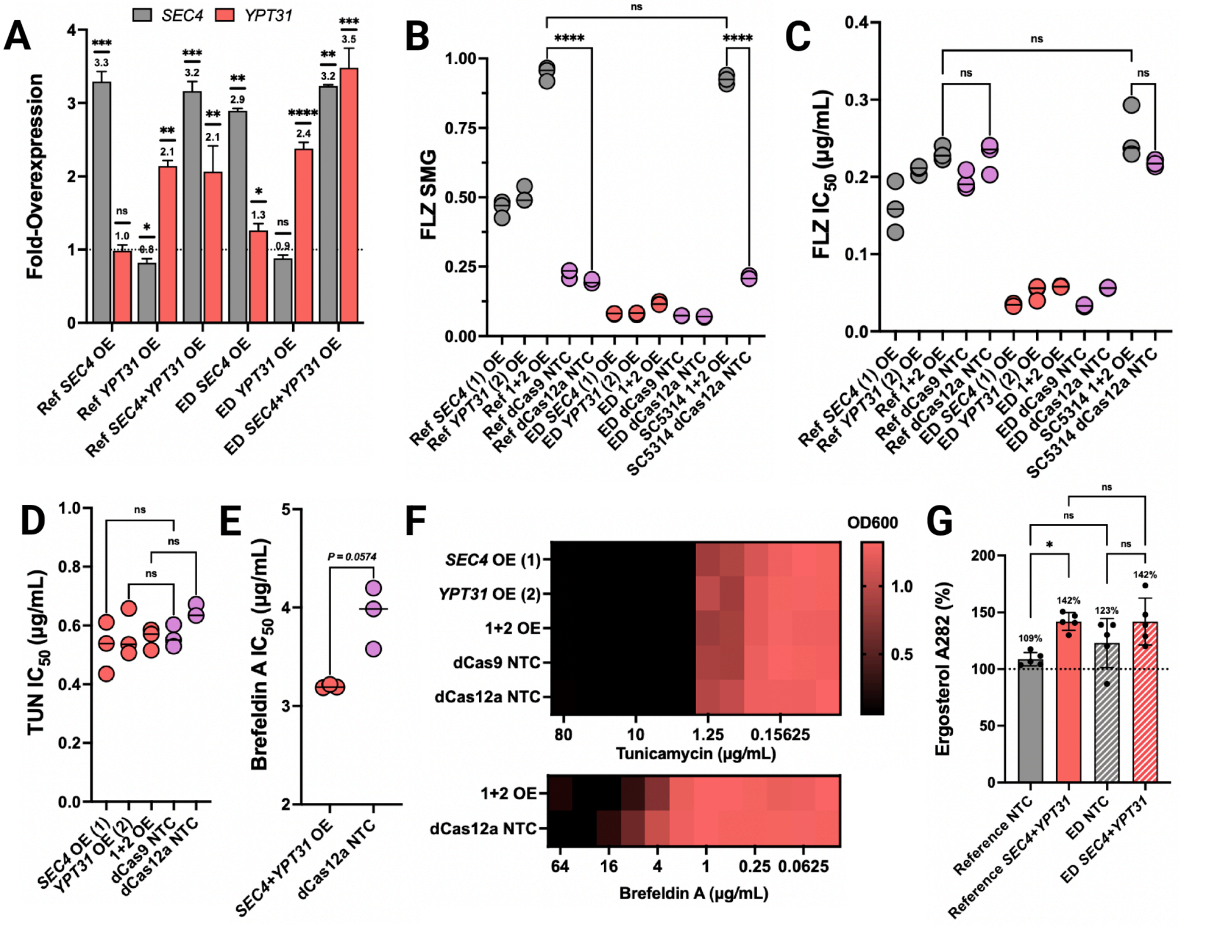


**Figure S11: Characterizing the influence of *SEC4* and *YPT31* upregulation on growth during ER stress as well as during growth with fluconazole with or without efflux.**

**(A)** The level of overexpression of *SEC4* and *YPT31* in the multiplexed CRISPRa strains in the parental (CAF2-1) and efflux-deficient backgrounds. Statistics were performed by comparing the dCt values between the given strain and its corresponding non-targeting control strain using a Welch's two-tailed t-test, **P* < .05, ***P* < 0.01, ****P* < 0.001, *****P* < 0.0001. **(B)** The FLZ SMG and **(C)** $IC_{50}$ values as a result of *SEC4* and/or *YPT31* overexpression in the reference strain (Ref) along with the derived efflux-deficient (ED) strain, and WT SC5314. There was no difference in FLZ $IC_{50}$ or SMG between the reference strain that the efflux-deficient mutant was derived from and SC5314. Statistics were performed using an ordinary one-way ANOVA with Tukey's multiple comparison test, ns $P > 0.05$, *****P* < 0.0001. **(D)** The $IC_{50}$ for each strain in tunicamycin. Statistics were performed using an ordinary one-way ANOVA with Tukey's multiple comparison test. Tests shown are the given strain to its corresponding non-targeting control strain (NTC), ns $P > 0.05$. **(E)** The $IC_{50}$ for each strain in brefeldin A. Statistics were performed using a Welch's two-tailed t-test. Exact p-value is shown for clarity. **(F)** Heat map showing the growth of each strain after 48 h at 37°C in the tunicamycin and brefeldin A MIC assays, displaying a lack of SMG. **(G)** The absorbance values at 282nm (representative of ergosterol abundance) for each strain reported as a % differential from the euploid WT NTC strain. Statistics were performed using an ordinary one-way ANOVA with Šídák's multiple comparisons test, ns $P > 0.5$, *$P < 0.05$.

**Part B: Supplementary Methods**

*Principal Component Analysis and PERMANOVA*

RNA-seq count data were first normalized by sequencing depth, and lowly expressed genes were removed. Principal component analysis (PCA) was performed on centered and scaled log-transformed counts-per-million (logCPM) values to visualize global transcriptional differences among samples. Variation due to FLZ treatment, genotype, and the corresponding FLZ/genotype interaction was quantified using permutational multivariate analysis of variance (PERMANOVA) with 999 permutations on the Euclidean distances between groups, calculated from the logCPM values. Differences in within-group variability were assessed to verify that the PERMANOVA results were not simply due to differences in dispersion within the groups. The code used to perform the PCA and PERMANOVA can be seen on our GitHub: https://github.com/TheShapiroLab/ChromosomeR.

*Multi-Omics Integration and Candidate Gene Selection*

To filter the list of significant sgRNAs from the CRISPRa screens, the 95% confidence interval of the non-targeting control sgRNAs in both concentrations of FLZ included log2 FCs of up to ~0.98. Therefore, we set a score of log2 FC of >1.5 in either FLZ concentration, or a log2 FC of >1.0 in both FLZ concentrations, as our threshold for significantly enriched sgRNAs in FLZ. As we have seen similarly in other CRISPR-dCas screens[3], we found the uncloned plasmid sequence had a significant enrichment in the 64μg/mL FLZ CRISPRa screen, suggesting there may be an on-target effect of the placeholder barcode in the empty vector.

Of the remaining 56 sgRNAs that passed the CRISPRa selection criteria, we further scored them based on whether they were upregulated in the ChrR-trisomic strains during growth in FLZ. We considered the differential expression of genes in both trisomic strains compared to the WT in FLZ, as opposed to the aneuploid-conditioned FLZ responses, since ChrR-mediated drug tolerance could be due to the influence of the trisomy alone and is not necessarily a result of the interaction between FLZ and ChrR aneuploidy, specifically. We further scored the list by whether the sgRNA was significantly depleted in either of the FLZ CRISPRi screens, and not depleted in the plain YPD CRISPRi screen, as we were specifically interested in whether the corresponding gene had a role in azole tolerance. Using the thresholds set by the non-targeting control sgRNAs in our CRISPRi screens, we initially identified that only 1/56 of the enriched CRISPRa sgRNAs were significantly depleted in either concentration of FLZ in the CRISPRi screens and not in plain YPD. Therefore, we relaxed the CRISPRi thresholds to a significant log2 FC of < -1.0 in either concentration of FLZ. This led to 4 sgRNAs that met all of the above CRISPRa, RNA-seq, and CRISPRi criteria. However, for these 4 sgRNAs, we additionally validated that they were indeed repressing their target gene via CRISPRi to confirm that they were not obviously false positives (**FIGURE S6C**). In addition to these 4 sgRNAs, 6 sgRNAs that had the next highest log2 FC in 64 μg/mL FLZ were selected, as well as another 4 sgRNAs that had the next highest log2 FC in 1 μg/mL FLZ, since some sgRNAs may not have led to successful repression with CRISPRi, and would therefore not have led to a significant change.

Reconstructing these 14 overexpression strains revealed that only one of them (targeting *CR_07920W_A*) did not show significant differential expression of the corresponding target gene. However, the sgRNA targeting *CR_09460C_A* was repressing its target (**FIGURE S6B**). Even though it has previously been shown that CRISPRa can cause gene repression and corresponding phenotypes[10], we were interested solely in overexpression-mediated drug tolerance. Therefore, we proceeded with the remaining 12 sgRNAs for all initial characterization assays.

*STRING Analysis*

STRING analysis was performed with the proteins corresponding to the significantly enriched CRISPRa sgRNAs in either FLZ concentration[11]. Two sgRNAs were excluded as they both targeted *CR_01870C*, which is a dubious ORF, and we were therefore unable to find an associated protein accession code[12]. However, the fact that multiple sgRNAs targeting this gene resulted in an enriched score in the CRISPRa screens in fluconazole suggests that further inquiry into this feature or potentially reclassifying it will reveal a previously unknown role in *C. albicans* drug response[13]. All of the results from the STRING analysis are available in **TABLE S1**.

*Clamped Homogeneous Electric Field (CHEF) Electrophoresis*

*C. albicans* strains SC5314, GC75, and CAY6440 were streaked onto YPD agar and grown for 24 h at 37°C. A single representative colony for each isolate was inoculated into 20 mL YPD and incubated at 30°C with 200 rpm shaking for 24 h. Subcultures were prepared by adding 50 µL of starter culture into fresh YPD medium, either containing 64 µg/mL FLZ or DMSO alone. FLZ flasks were prepared in duplicate for each isolate. Cultures were incubated at 37°C with 200 rpm shaking for 24 h. Five mL of each culture were subcultured into a fresh flask of the same composition (FLZ or DMSO alone) and the subcultures incubated for another 24 h at 37°C with shaking. Cells were harvested and embedded in agarose to make plugs for CHEF analysis as described previously[14]. CHEF gels were run at 12°C, 2.5 V/cm, with a 120-degree angle. Initial conditions used 24 h with a 120-300 sec linear ramp followed by 46 h with a 420-900 s linear ramp. A longer run time was also used to attempt to separate the largest chromosomes (45 h with a 180-300 s linear ramp followed by 99 h with a 420-900 s linear ramp). Gels were stained using ethidium bromide, then destained in 1x TAE buffer before imaging.

### <u>Part C: Description of Supplementary Tables</u>

1. TABLE S1: Screening Data
2. TABLE S2: Library Construction
3. TABLE S3: Growth Profiling Data
4. TABLE S4: Primer/gRNA/Strain Log
5. TABLE S5: RT-qPCR and ddPCR Data

### Supplementary References


1. Gervais, N. C. *et al.* Development and applications of a CRISPR activation system for facile genetic overexpression in Candida albicans. *G3 (Bethesda)* **13**, (2023).
2. Wensing, L. *et al.* A CRISPR interference platform for efficient genetic repression in Candida albicans. *mSphere* **4**, (2019).
3. Wensing, L. F. *et al.* Pooled CRISPRi screening reveals fungal-specific drug target candidates. *Nat. Microbiol.* 1–13 (2026).
4. Delarze, E. *et al.* Identification and characterization of mediators of fluconazole tolerance in candida albicans. *Front. Microbiol.* **11**, 591140 (2020).
5. Kukurudz, R. J. *et al.* Acquisition of cross-azole tolerance and aneuploidy in Candida albicans strains evolved to posaconazole. *G3 (Bethesda)* **12**, (2022).
6. Gautier, C. *et al.* Sphingolipid Homeostasis, Mitochondrial Activity, and PKA Signaling Drive an Azole-Tolerant State. *Microbiology* (2025).
7. Szklarczyk, D. *et al.* The STRING database in 2025: protein networks with directionality of regulation. *Nucleic Acids Res.* **53**, D730–D737 (2025).
8. Rudner, A. D. & Murray, A. W. Phosphorylation by Cdc28 activates the Cdc20-dependent activity of the anaphase-promoting complex. *J. Cell Biol.* **149**, 1377–1390 (2000).
9. Bishop, A. *et al.* Hyphal growth in Candida albicans requires the phosphorylation of Sec2 by the Cdc28-Ccn1/Hgc1 kinase. *EMBO J.* **29**, 2930–2942 (2010).
10. Maroc, L., Shaker, H. & Shapiro, R. S. Functional genetic characterization of stress tolerance and biofilm formation in Nakaseomyces (Candida) glabrata via a novel CRISPR activation system. *mSphere* **9**, e0076123 (2024).
11. Szklarczyk, D. *et al.* The STRING database in 2023: protein-protein association networks and functional enrichment analyses for any sequenced genome of interest. *Nucleic Acids Res.* **51**, D638–D646 (2023).
12. Lew-Smith, J., Binkley, J. & Sherlock, G. The Candida Genome Database: annotation and visualization updates. *Genetics* **229**, iyaf001 (2025).
13. Li, Q.-R. *et al.* Revisiting the Saccharomyces cerevisiae predicted ORFeome. *Genome Res.* **18**, 1294–1303 (2008).
14. Hoyer, L. Visualization of Yeast Chromosomes Using Clamped Homogeneous Electric Field (CHEF) Electrophoresis. *protocols.io* https://www.protocols.io/view/visualization-of-yeast-chromosomes-using-clamped-h-8epv5jdpdl1b/v1 (2023).
